# Increasing brain half-life of antibodies by additional binding to myelin oligodendrocyte glycoprotein, a CNS specific protein

**DOI:** 10.1101/2024.11.04.621218

**Authors:** Marie-Lynn Cuypers, Tom Jaspers, Jarne Clerckx, Simon Leekens, Christopher Cawthorne, Guy Bormans, Frederik Cleeren, Nick Geukens, Bart De Strooper, Maarten Dewilde

## Abstract

**Background:** Therapeutic antibodies for the treatment of neurological disease show great potential, but their applications are rather limited due to limited brain exposure. The most well-studied approach to enhance brain influx of protein therapeutics, is receptor-mediated transcytosis (RMT) by targeting nutrient receptors to shuttle protein therapeutics over the blood-brain barrier (BBB) along with their endogenous cargos. While higher brain exposure is achieved with RMT, the timeframe is short due to rather fast brain clearance. Therefore, we aim to increase the brain half-life of antibodies by binding to myelin oligodendrocyte glycoprotein (MOG), a CNS specific protein.

**Methods:** Alpaca immunization with mouse/human MOG, and subsequent phage selections and screenings for MOG binding VHHs were performed to find mouse/human cross-reactive VHHs. Their ability to increase the brain half-life of antibodies was evaluated by coupling two different MOG VHHs (low/high affinity) in a mono- and bivalent format to an anti-β-secretase 1 or anti-SarsCov2 antibody fused to an anti-transferrin receptor (TfR) VHH for active transport over the BBB. Brain pharmacokinetics and pharmacodynamics, CNS and peripheral biodistribution, and brain toxicity were evaluated after intravenous administration to balb/c mice.

**Results:** Additional binding to MOG increases the C_max_ and brain half-life of antibodies that are actively shuttled over the BBB. biMOG_Low_:monoTfR:SarsCov antibodies could be detected in brain 49 days after a single intravenous injection, which is a major improvement compared to monoTfR:SarsCov antibodies which cannot be detected in brain anymore one week post treatment. Additional MOG binding of antibodies does not affect peripheral biodistribution but alters brain distribution to white matter localization and less neuronal internalization.

**Conclusions:** We have discovered mouse/human/cyno cross-reactive anti-MOG VHHs which have the ability to drastically increase brain exposure of antibodies. Combining MOG and TfR binding leads to distinct PK, biodistribution, and brain exposure, differentiating it from the highly investigated TfR- shuttling. It is the first time such long brain antibody exposure is demonstrated after one single dose.

This new approach of adding a binding moiety for brain specific targets to RMT shuttling antibodies is a huge advancement for the field and paves the way for further research into brain half-life extension.

## BACKGROUND

Central nervous system (CNS) disorders, including Alzheimer’s disease, brain cancer, multiple sclerosis, and Parkinson’s disease, are among the most prevalent yet poorly treatable illnesses, significantly impacting quality of life and placing an economic burden on healthcare systems. Neurological disorders, responsible for 9 million annual deaths, rank as the second primary cause of mortality (1). The aging global population is expected to further escalate these numbers, with one in three people projected to develop a neurological disorder (1). Over the past five years, the CNS pipeline has expanded 31 %, constituting 14 % of the total industry research and development (R&D) pipeline, making it the second-largest therapy area, following oncology (2,3).

Developing effective therapies for CNS disorders faces significant challenges due to the elusive blood-brain barrier (BBB), a selectively permeable structure, separating the brain parenchyma from the circulation. While important to protect the brain, this barrier also limits the entry of the majority of drug compounds into the brain increasing the failure rate of CNS drug discovery. The primary component of the BBB is a tight-junction complex of non-fenestrated brain microvascular endothelial cells, supported by astrocytes, pericytes, and two basement membranes (4). Only 0.1 % of circulating antibodies are estimated to reach the brain at steady-state concentration, necessitating high dose administrations with potential peripheral side effects (5). Given the severe consequences and the increasing prevalence of major CNS-related diseases, such as Alzheimer’s disease or Parkinson’s disease, there is an urgent need for novel and effective treatment options that can overcome the BBB.

In this respect, invasive methods such as direct intracerebral delivery or BBB disruption are explored, but they are less suitable for chronic treatments due to patient discomfort and infection risks (6). In the past years, extensive R&D efforts have yielded innovative BBB penetration technologies, with a notable non-invasive approach using monoclonal antibodies (mAb) or fragments thereof coupled to a therapeutic cargo targeting nutrient receptors that shuttle the BBB (e.g. transferrin receptor (TfR), insulin receptor, CD98, …), facilitating receptor-mediated transcytosis (RMT) of therapeutic cargos into the brain (6–11). Multiple drug candidates are actively being investigated in the clinic, and a first drug candidate reaching the brain through RMT has been approved in 2021 in Japan (Izcargo®) (12–14). In addition, recently two antibodies targeting Aβ in brains of patients with Alzheimer’s disease were approved for clinical use by the FDA (15). These antibodies were not engineered to actively cross the BBB but rely on sufficient passive diffusion over the BBB and binding to brain enriched targets. Inevitably, this strategy relies on relatively high peripheral dosing with an increased risk of dose related side effects, like ARIA (amyloid-related imaging abnormalities), resulting in brain swelling and bleeding with a potential lethal outcome (15,16). Therefore, the field is heavily investigating methods to increase antibody yields specifically in the brain, also for the anti-Aβ antibodies as exemplified by the clinical trials of Roche with trontinemab, which is basically Roche’s anti-Aβ mAb (gantenerumab) fused to an anti-TfR mAb to allow RMT to the brain (NCT04639050).

Several studies have shown the success of this noninvasive TfR BBB shuttling technology to increase brain concentrations of therapeutics (7,8,12,17–19). However, the main downside of using TfR for RMT, is the widespread expression of TfR in multiple peripheral organs, leading to fast peripheral clearance and hence lower central exposure. Therefore, the observed therapeutic effect is rather short and in a clinical setting, more frequent intravenous dosing might be required to keep levels high enough to elicit a therapeutic effect.

More recently, an alternative possibility to decrease the brain efflux of antibodies after their limited passive diffusion into the brain was explored (20). To decrease its efflux, a therapeutic protein was fused to a mAb fragment targeting an abundant brain protein, i.e. myelin oligodendrocyte glycoprotein (MOG), which resulted in significantly increased brain accumulation of the antibody-drug conjugate. However, the achieved brain concentrations remained rather low, and this research group did not show any prolonged therapeutic effect (20,21).

Here we report the development of mouse/human/cyno cross-reactive anti-MOG single variable domain antibodies (VHHs or nanobodies). First, we’ve evaluated the role of the affinity and valency of these anti-MOG VHHs fused to an antibody on the levels and activity of the fused antibody, this both in plasma and brain. Next, we studied in depth the systemic clearance, brain pharmacokinetics (PK) and pharmacodynamics (PD), CNS and peripheral biodistribution, and brain toxicity for the selected lead molecule. We demonstrate that MOG binding VHHs have the ability to dramatically increase the CNS half-life of two different antibodies that are actively shuttled over the BBB, and to significantly prolong the therapeutic effect of an anti-β-secretase 1 (BACE1)/anti-TfR antibody construct. This new approach of combining brain specific target binding and RMT shuttling of antibodies marks a significant advancement in the pursuit of brain accumulation of biologicals, addressing a major bottleneck in the search for innovative drugs for neurological disorders.

## MATERIAL & METHODS

### VHH Library Generation

Three alpacas were subjected to three rounds of four bi-weekly DNA immunizations (VIB Nanobody Core, Brussels, Belgium), at a one-month interval, using a pool of recombinant pVAX1 plasmids encoding various proteins, including human myelin oligodendrocyte glycoprotein (hMOG). Seven months later, the same animals were immunized with another pool of recombinant pVAX1 plasmid DNA encoding various proteins, including mouse MOG (mMOG). The immunization with the mouse equivalent consisted of two rounds, round one consisting of one and round two of 3 bi-weekly DNA immunizations. DNA solutions were injected intradermally at multiple sites at front and back limbs near the draining lymph nodes followed by electroporation. A total amount of 2 mg of each pool of the plasmid DNA was used per injection per animal. Four and 8 days after the first and the last mMOG immunization, 100 mL anticoagulated blood was collected from each animal for the preparation of peripheral blood lymphocytes. The VHH encoding genes were recovered and the phagemid library was cloned as previously prescribed (22). In short, total RNA from peripheral blood lymphocytes was used as template for first strand cDNA synthesis with oligodT primers. This cDNA was used to amplify the VHH-encoding open reading frames by PCR, digested with PstI and NotI, and cloned into a phagemid vector. The library was transformed into electro-competent *E.coli* TG1 cells, which resulted in 10^8^ independent transformants, of which at least 85 % contained the vector with the right insert size.

### Cell line generation

Stable Chinese hamster ovary (CHO) cell lines, overexpressing mouse, human or cynomolgus MOG (cMOG), were generated with the Flp-In^TM^ CHO system (Invitrogen, Waltham, MA, USA). DNA encoding m/h/cMOG, tagged with c-terminal hemagglutinin followed by green fluorescent protein (GFP) under the control of an internal ribosome entry site, was synthesized by Twist Bioscience (San Francisco, CA, USA) and subcloned into the pcDNA5/FRT mammalian expression vector (Invitrogen, Waltham, MA, USA). This target expressing vector and the Flp-In^TM^ recombinase vector pOG44, were co-transfected in the Flp-In^TM^ CHO cell line using Xtreme gene HP DNA transfection reagent (Sigma, St. Louis, MO, USA). Cells were cultured in Ham’s F-12 Nutrient Mix medium supplemented with GlutaMAX (Gibco, Thermo Fisher Scientific, Waltham, MA, USA), 10 % fetal bovine serum (FBS) (Gibco, Thermo Fisher Scientific, Waltham, MA, USA) and 100 µg/mL zeocin selection antibiotic (Invivogen, San Diego, CA, USA) before transfection or 700 µg/mL Hygromycin B gold (Invivogen, San Diego, CA, USA) after transfection to select for stably transfected cells. Stable cell lines were amplified and aliquots frozen with 10 % dimethylsulfoxide (DMSO) (Sigma, St. Louis, MO, USA) for further use.

### Isolation of anti-MOG VHHs

To select species cross-reactive anti-MOG VHHs, two rounds of cell selections on 5×10^6^ mMOG expressing cells (in-house cells derived from Flp-In^TM^ CHO Cell Line, Invitrogen, Waltham, MA, USA) were performed followed by one round of in solution selection with 50 nM in-house biotinylated human MOG (hMOG) (R&D Systems, Minneapolis, MN, USA). Thereafter, the library was subcloned into an expression vector (pBDS100, a modified pHEN6 vector with an OmpA signal peptide and a C-terminal 3xFlag/6xHis tag). The expression library was used to transform TG1 *E.coli* after which VHHs were expressed from single colonies. These VHHs were screened for direct binding to the biotinylated hMOG using the AlphaScreen FLAG (M2) Detection Kit (6760613, PerkinElmer, Waltham, MA, USA) and to the mMOG overexpressing CHO cells using flow cytometry with mouse-anti-FLAG-iFluor647 Ab (Genscript, Piscataway, NJ, USA). The cross-reactive hits were sequenced and clustered according to sequence homology. One representative of each cluster was expressed following the protocol by Pardon et al. (22) and purified using AmMag^TM^ Ni-charged magnetic beads and an AmMag^TM^ SA Plus System (Genscript, Piscataway, NJ, USA).

### Monoclonal antibodies

The control antibody consisted of either the anti–BACE1 Fab (1A11) (23) or an anti-SARS-CoV-2 spike protein Fab (24), and Nb62 binding to mTfR (7) or an anti-GFP VHH, fused to a human IgG1 Fc domain with LALA-PG effector and complement knock-out mutations (L234A, L235A, P329G) and knob-into-hole mutations (T350V, T366L, K392L, T394W/ T350V, L351Y, F405A, Y407V) to generate heterodimeric antibodies (monoTfR:BACE1 and monoTfR:SarsCov). MOG binding antibodies were created by genetically fusing these controls with either one (monovalent) or two (bivalent) MOG binding VHHs with different affinities (BBB00498 and BBB00500) at the c-terminal side of the Fc (Fig. 1). Encoding genes were ordered at Twist Bioscience (San Francisco, CA, USA) in pTwist CMV BG WPRE Neo vector (Twist Bioscience, San Francisco, CA, USA). Antibodies were expressed in expiCHO-S^TM^ cells (Thermo Fisher Scientific, Waltham, MA, USA) using CHOgro high yield expression system (Mirus Bio, Madison, WI, USA) according to the manufacturer’s protocol and purified following the protocol by Nesspor et al. (25). The purification protocol consisted of protein A purification using AmMag^TM^ Protein A Magnetic Beads (Genscript, Piscataway, NJ, USA), followed by purification over a CaptureSelect™ CH1-XL Pre-packed Column (Thermo Fischer Scientific, Waltham, MA, USA) and size exclusion chromatography on a Superdex 200 Increase 10/300 GL (Cytiva, Marblorough, MA, USA). Generated antibodies were monoTfR:BACE1, monoMOG_Low_:monoTfR:BACE1, monoMOG_High_:monoTfR:BACE1, biMOG_Low_:monoTfR:BACE1, biMOG_High_:monoTfR:BACE1, monoGFP:BACE1, biMOG_Low_:monoGFP:BACE1, monoTfR:SarsCov, biMOG_Low_:monoTfR:SarsCov, monoGFP:SarsCov, biMOG_Low_:monoGFP:SarsCov (Fig. 1).

**Figure 1.**
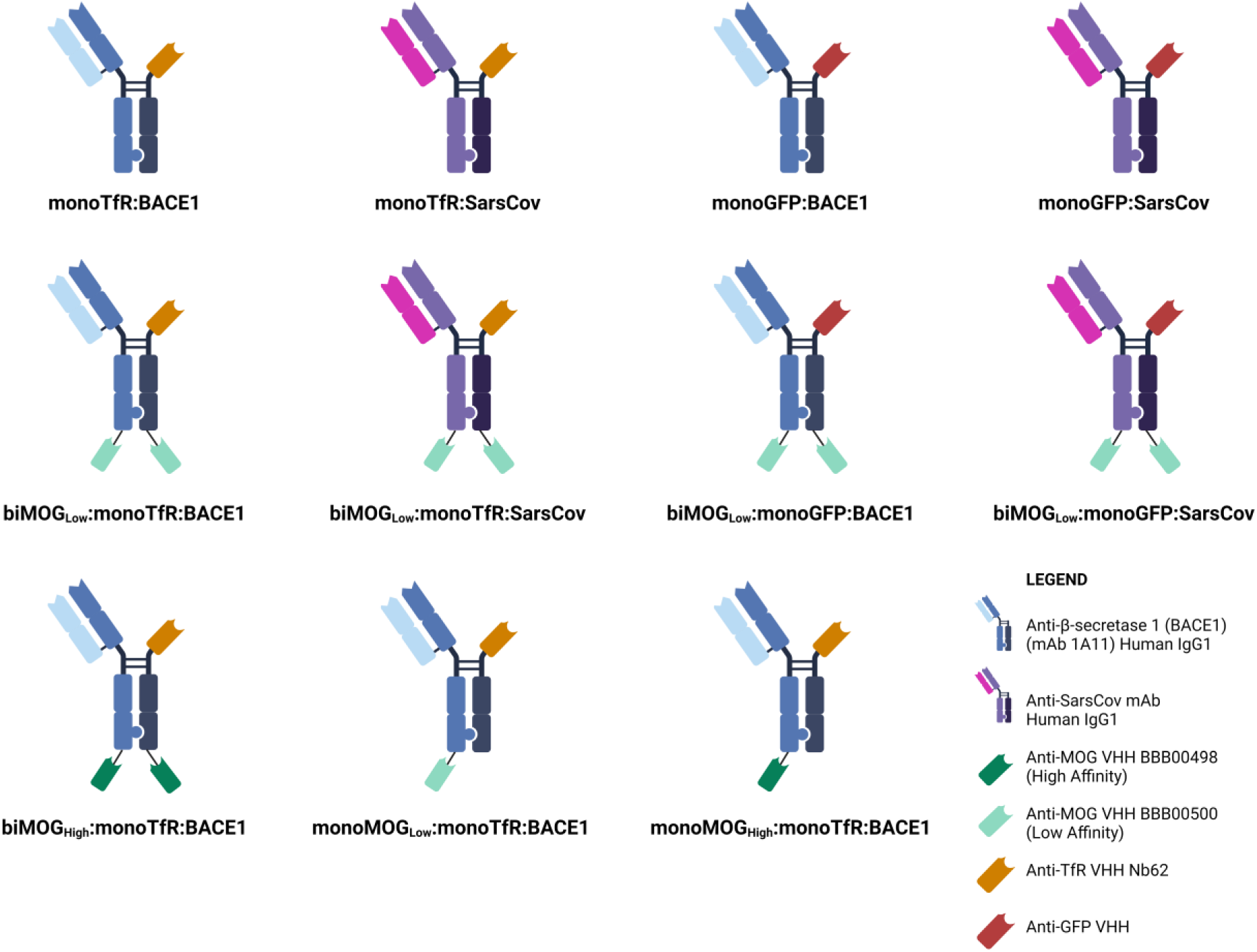
Multispecific antibody constructs. Schematic overview of produced antibodies binding to BACE1 or SarsCov, TfR or GFP, and MOG.

### In vitro binding and affinity determination

Binding of purified VHHs or antibodies to m/h/cMOG expressing cells was evaluated by flow cytometry. 10^5^ MOG/GFP expressing CHO cells were incubated, 30 min on ice, with the purified proteins in a concentration range between 0.05 and 5000 nM. Cells were washed twice with PBS containing 2 % FBS (Gibco, Thermo Fisher Scientific, Waltham, MA, USA). After washing, cells were stained, 30 min on ice in the dark, with mouse-anti-FLAG-iFluor647 (1:500) (Genscript, Piscataway, NJ, USA) or anti-human IgG Fc Alexa Fluor 647 (1:200) (Thermo Fisher Scientific, Waltham, MA, USA), for VHHs or antibodies respectively, and eBioscience Fixable Viability Dye eFluor 780 (1:2000) (Invitrogen, Waltham, MA, USA) in staining buffer (PBS + 2 % FBS), for protein detection and live/dead staining respectively. After staining, cells were washed with staining buffer, followed by fixation with 4 % paraformaldehyde (PFA) (Alfa Aesar, Haverhill, MA, USA) (15 min, room temperature (RT), protected from light), and resuspension in staining buffer. Cells were kept at 4 °C before analysis. Attune NxT Flow Cytometer (Thermo Fisher Scientific, Waltham, MA, USA) was used for sample analysis. CHO-GFP expressing cells were used for background correction. Data analysis was performed using FCS Express 7 software (De Novo Software, Pasadena, CA, USA).

### Animals

Male and female Balb/c mice (Balb/cAnNRj), 7-8 weeks old, with an approximate weight of 18–24 g were purchased at Janvier (Janvier, Le Genest-Saint-Isle, France). All animal experiments were approved by the KU Leuven Animal Ethics Committee (project P091/2022 and P038/2024) following governmental and EU guidelines. Mice were housed under a 12-h light/dark cycle with water and standard rodent diet ad libitum.

### Intravenous injection

Intravenous injections of antibodies were performed by putting the mice in a heating chamber at 40 °C for 10 minutes. Afterwards, mice were put in a restrainer and mAbs were injected in the tail vein at volumes between 45 and 120 µL.

### Plasma/brain sampling

Blood collection was performed by heart puncture at time of brain collection in K3 EDTA coated tubes (Sarstedt, Nümbrecht, Germany). Blood was processed to plasma by centrifugation (10 min, 2000*g*, 4 °C and 10 min, 16 000*g*, 4 °C), and stored at −20 °C. Mice were euthanized with a Dolethal® (pentobarbital) (Vetoquinol, Niel, Belgium) overdose (150–200 mg/kg) injected peritoneally. Brains were harvested after transcardial perfusion with heparinized (Leo Pharma, Ballerup, Denmark) PBS, snap frozen by submerging collection tubes in liquid nitrogen and stored at −80 °C until further processing.

### Aβ1-40 detection using MSD ELISA

A brain hemisphere was homogenized in buffer containing 0.4 % diethylamine (Sigma, St. Louis, MO, USA) and 50 mM NaCl (Fisher Scientific, Waltham, MA, USA), supplemented with cOmplete™ protease inhibitor cocktail EDTA-free (Roche Diagnostics, Mannheim, Germany) using Ceramic Bead Lysing Matrix D 1.4 mm (MP Biomedicals, Irvine, CA, USA) and a FastPrep-24 classic homogenizer (MP Biomedicals, Irvine, CA, USA)(6 m/s, 3x 20 seconds, 5 min cooling on ice between different cycles). Homogenate was subjected to high speed centrifugation (100 000*g*, 1 h, 4 °C) and supernatant was neutralized with 0.5M Tris-HCl (pH 6.8) (Sigma, St.Louis, MO, USA). Aβ_1-40_ levels in brain and plasma samples were quantified by ELISA using Meso Scale Discovery (MSD, Meso Scale Diagnostics, Rockville, MD, USA) 96-well plates and in-house produced antibodies. MAb LTDA_Aβ40, which recognizes the C-terminus of Aβ_1-40,_ was used as a capture antibody and LTDA_hAβN, a rodent sulfoTAG-fused antibody recognizing the N-terminus as the detection antibody. MSD plates were coated overnight (ON) at 4 °C with mAb LTDA_Aβ40 (1.5 µg/mL in PBS, 50 µL/well). Next day, plates were washed 3x with PBS-T (0.05 % Tween 20, Sigma, St. Louis, MO, USA) and blocked with PBS 0.1 % casein (Sigma, St. Louis, MO, USA) (150 µL/well) for 4 hours at RT, shaking 700 rpm. Plates were washed 3x with PBS-T and incubated with samples (50 µL/well) ON at 4 °C. Sample mix consisted of plasma or 1:2 diluted brain homogenate in PBS 0.1 % casein, diluted 1:2 with LTDA_hAβN sulfoTAG (1:2000 in PBS/casein), pre-incubated 15 min at RT, shaking 700 rpm. Serial two-fold dilutions of rodent Aβ_40_, with concentrations ranging between 2.4 and 2500 pg/mL, were used as calibration curve. Next day, plates were washed 3x with PBS-T and signal measured with MSD plate reader after adding 150 µL of READ buffer (R92TC-1 diluted 1:2 in water, Meso Scale Diagnostics, Rockville, MD, USA). Sample concentrations were calculated based on the Aβ_40_ calibration curve using a nonlinear fit, Log(agonist) vs response – variable slope (4 parameters) (GraphPad Prism 10, GraphPad Software, San Diego, CA, USA).

### Antibody detection in brain and plasma

The antibody levels in plasma and brain homogenate were quantified using an in-house developed ELISA. A brain hemisphere was homogenized with 9 volumes of PBS containing 1 % Igepal (Alfa Aesar, Haverhill, Massachusetts, USA), supplemented with cOmplete™ protease inhibitor cocktail EDTA-free (Roche Diagnostics, Mannheim, Germany) using Ceramic Bead Lysing Matrix D 1.4 mm (MP Biomedicals, Irvine, CA, USA) and a FastPrep-24 classic homogenizer (MP Biomedicals, Irvine, CA, USA) (6.5 m/s, 2x 45 seconds, 1 min cooling on ice between different cycles). After a 45 min incubation on ice, homogenate was centrifuged (16 000*g*, 10 min, 4°C) and supernatant collected. This extraction procedure was repeated two additional times with the obtained pellet. Plates were coated ON at 4 °C with 2 µg/mL mouse anti-human IgG Fc (Genscript, Piscataway, NJ, USA) in PBS. Plates were blocked with PBS 0.1 % casein (Sigma, St. Louis, Missouri, USA) for two hours at RT. Then, samples were diluted in PBS 0.1 % casein, added to the plate and incubated for two hours at RT under shaking conditions (300 rpm). Serial two-fold dilutions of the purified antibody constructs in corresponding matrix (diluted plasma or brain homogenate), with concentrations ranging between 16 and 0.125 ng/mL, were used as calibration curves. Detection of the captured human IgGs was performed with mouse anti-human IgG Fab-HRP (Genscript, Piscataway, NJ, USA) (1:5000 dilution in PBS 0.1 % casein). Each incubation step, except blocking, was preceded by a washing step with PBS 0.002 % Tween 80. Plates were developed for 30 minutes using 1-step Ultra TMB-ELISA substrate solution (Thermo Fisher Scientific, Waltham, MA, USA). The reaction was stopped with 2 M H_2_SO_4_. Absorption was measured at 450 nm using an ELx808 Absorbance Microplate Reader (BioTek Instruments, Bad Friedrichshall, Germany). IgG concentrations in the samples were calculated based on the human IgG calibration curve using a linear regression fit (GraphPad Prism 10, GraphPad Software, San Diego, California, USA).

### Brain immunohistochemistry

After perfusion with PBS, mouse brains were fixed ON in 4 % PFA (Alfa Aesar, Haverhill, MA, USA) (4 °C), followed by storage in PBS/0.2 % Sodium azide (Sigma, St. Louis, MO, USA) (4°C) for maximum two weeks. Brains were embedded in 4 % agar in PBS on the day of sectioning. Sagittal brain sections (30 µm) were cut using a Leica VT1000S vibratome (Leica, Wetzlar, Germany) and stored in PBS/0.2 % Sodium azide until further processing. Tissue slices were blocked and permeabilized with 5 % BSA (Sigma, St. Louis, MO, USA), 0.3 % Triton-X 100 (Sigma, St. Louis, MO, USA) in PBS (1.5h, RT), followed by fluorescent staining with primary (ON, 4°C) and secondary antibodies (2h, RT) (Table S1) diluted in PBS containing 1 % BSA, 0.3 % Triton-X 100. Sections were mounted on Superfrost plus slides (Fisher Scientific, Waltham, MA, USA) using Prolong gold anti fade mounting medium (Thermo Fisher Scientific, Waltham, MA, USA). Images were collected by a Zeiss Axioscan 7 or Zeiss LSM 880 confocal microscope with airyscan using a 63x oil-immersion objective (Zeiss, Oberkochen, Germany).

### Assessment of biodistribution using radiolabeled antibodies

Antibody constructs (5-10 mg/mL in PBS, pH 7.4) were buffer-exchanged to a metal-free NaHCO_3_ buffer (50 mM, pH 8.5) via SE-HPLC using a Superdex200 Increase 10/300 GL column (Cytiva, Marlborough, MA, USA) at a flow rate of 0.75 mL/min, executed on an Agilent 1100 Series HPLC Value System, with a variable wavelength detector (Agilent, Santa Clara, CA, USA) set to 280 nm. Following buffer exchange, a mixture of the antibody constructs with a 10-fold molar excess of DFO*-NCS (5 mg/mL in DMSO) (ABX, Radeberg, Germany) was incubated ON at 4 °C with gentle shaking (Eppendorf ThermoMixer, Merck, Darmstadt, Germany). The DFO*-derivatized constructs were purified by SE-HPLC under the same conditions as described above.

DFO*-antibody constructs were labeled with zirconium-89 (^89^Zr; t_1/2_ = 78.41 h) (BV Cyclotron UV, Amsterdam, the Netherlands), a radionuclide commonly employed for positron emission tomography (PET). Radiolabeling was conducted under mild conditions (0.25 M HEPES buffer, pH 7.2, 37 °C, gentle shaking). The crude radiolabeling mixture was purified through successive cycles of ultracentrifugation using Amicon Ultra-4 centrifugal filter units (Merck, Darmstadt, Germany, MWCO 30 kDa) with the addition of formulation buffer (5 mg/mL ascorbic acid in PBS, pH 7.4). Radiochemical purity was assessed quantitatively via radio-SE-HPLC as described above with radiometric detection (GABI* radioactivity HPLC flow detector, Elysia-Raytest, Liège, Belgium).

The stability of the radiolabeled antibody constructs was assessed over a period of 7 days in two different conditions: formulation buffer at 4 °C and normal mouse serum at 37 °C. Stability was determined via radio-instant thin layer chromatography (radio-iTLC) using iTLC-SG paper (Glass microfiber chromatography paper impregnated with silica gel, Agilent Technologies, Folsom, CA, USA), developed in a citric acid solution (20 mM, pH 5), imaged via autoradiography using phosphor storage screens (super-resolution screen, PerkinElmer, Waltham, MA, USA), and analyzed by Cyclone Plus system (PerkinElmer, Waltham, MA, USA) and OptiQuant software (PerkinElmer, Waltham, MA, USA). Stability was determined as the percentage of total activity corresponding to protein-associated activity, identified by an Rf-value of 0.

Mice were administered ^89^Zr-DFO*-antibody constructs via tail vein injection, (1, 2, 4, or 6 MBq for groups sacrificed at 1, 3, 7, or 14 days, respectively). The molar amount of injected antibody construct ranged between 174-362 nmol/kg.

Static PET images (15 min) were acquired for 4 out of 5 animals before sacrifice using a β-cube PET scanner (Molecubes, Ghent, Belgium). Mice were anesthetized (5 % isoflurane in O_2_ at 1 L/min flow rate for induction, 2.5 % for maintenance) and placed into the imaging cell (Molecubes, Ghent, Belgium) before transfer to the scanner. Temperature and respiration were monitored throughout the scan. Following PET scanning, computed tomography (CT) imaging was performed for anatomical co-registration using an X-cube CT scanner (Molecubes) using the ‘General Purpose’ protocol with the following parameters: 50 kVp, 480 exposures, 85 ms/projection, 100 μA tube current, rotation time 60 s.

PET and CT images were fused and the PET image scaled to standardized uptake value (SUV). Uptake in brain, liver, muscle and heart region (used as blood surrogate) was quantified via regions-of-interest drawn on CT data. Coronal maximum intensity projections of the PET data were generated to illustrate tracer distribution at each timepoint. All image analysis was done using PFUS v4.0 (PMOD Technologies GmbH, Zurich, Switzerland). Following PET/CT imaging, mice were euthanized with a Dolethal® (200 mg/mL pentobarbital) (Vetoquinol, Niel, Belgium) overdose (350 mg/kg) injected intraperitoneally. After transcardial perfusion with 0.9% NaCl, major organs were collected, weighed, and analyzed using a Wizard² Gamma Counter (2480-0010, PerkinElmer, Waltham, MA, USA) for biodistribution studies. Results are reported in SUV, which normalizes the tracer uptake for injected dose, organ/tissue weight, and body weight. Finally, mouse brains were rapidly frozen in 2-methylbutane cooled to approximately −35 °C. Horizontal sections of 20 µm were prepared using a Shandon cryotome FSE (ThermoFisher, Waltham, MA, USA), mounted on SuperFrost Plus adhesive microscope slides (ThermoFisher, Waltham, MA, USA), and imaged by autoradiography as previously described.

### MOG/BACE1/TfR quantification in brain using western blot

Total protein concentration of brain homogenates used for PK analysis was measured using a Pierce BCA Protein Assay kit (Thermo Fisher Scientific, Waltham, MA, USA). Samples containing 7,5 µg of total protein in NuPAGE LDS sample buffer (1x) (Thermo Fisher Scientific, Waltham, MA, USA) supplemented with 25mM DTT (reducing agent) (Thermo Fisher Scientific, Waltham, MA, USA), were heated for 5 minutes at 95°C, loaded onto NuPAGE 4-12 % Bis-Tris gels (Thermo Fisher Scientific, Waltham, MA, USA) and run 45 min at 200 V in MES buffer (Thermo Fisher Scientific, Waltham, MA, USA). Gels were transferred onto nitrocellulose membranes (0.45µm) (Cytiva, Amersham Biosciences, Amersham, UK) using a mini blot module (Thermo Fisher Scientific, Waltham, MA, USA) for wet transfer, followed by membrane blocking (5 % milk (Carl Roth, Karlsruhe, Germany) , 5 % BSA (Sigma, St. Louis, MO, USA), 0,1 % Tween 20 (Sigma, St. Louis, MO, USA) in PBS)(1h, RT) and primary (4°C, ON) and secondary (1h, RT) antibody incubations (Table S2). Mouse anti-β-actin HRP (Sigma, St. Louis, MO, USA) (A3854, 1:40 000, 30 min, RT) served as loading control. ECL Select western blotting reagent (Cytiva, Amersham Biosciences, Amersham, UK) and Chemidoc MP Imaging system (Bio-rad, Hercules, CA, USA) were used for detection. Analysis was performed with ImageLab 6.1 software (Bio-rad, Hercules, CA, USA).

### Statistical Analysis

Statistical analysis was performed using GraphPad Prism 10 (GraphPad Software, San Diego, CA, USA). Parametric methods were used for analysis as data distribution was sufficiently normal for all experiments. One-way ANOVAs (variable time) with Dunnett’s test for multiple comparisons were used for western blot analysis. For other experiments two-way ANOVAs (variables time and treatment) were performed. The reported F statistic and associated p-value were derived on the interaction term for comparison of PK/PD curves or on Dunnett’s multiple comparisons test for comparing single timepoint data to a control group. Table S3 and S4 include results for all one- and two-way ANOVA tests, and pairwise comparisons respectively.

## RESULTS

### Generation and expression of mouse/human cross-reactive MOG binding VHHs

To find mouse/human cross-reactive anti-MOG VHHs, a VHH-displaying phage library originating from camelids immunized with hMOG and mMOG, was subjected to one or two *in vitro* selection rounds on CHO-mMOG overexpressing cell lines and optional one selection round on recombinant hMOG to increase the likelihood of finding cross-reactive VHHs. Single clones from libraries with promising enrichment factors were sequenced and clustered according to homology. 93 unique clones were selected, expressed and periplasmic extracts were prepared. Screening results revealed 59 mMOG binders and 54 hMOG binders of which 24 VHHs were mouse/human cross-reactive. One mouse specific binder (BBB00497) and 5 cross-reactive VHHs (BBB00498-BBB00502) were selected based on sequence diversity and characteristics (framework identity to germline and post translational modification sites). These were recloned in an expression vector harboring a c-terminal 3xFLAG and a hexahistidine tag. VHHs were expressed in TG1 and purified from the periplasmic extract with Ni-charged AmMag^TM^ beads.

### In vitro characterization of anti-MOG VHHs

*In vitro* binding characteristics of these six VHHs were determined with flow cytometry on mouse, human and cynomolgus MOG overexpressing cell lines using mouse-anti-FLAG-iFluor647 detection antibody. Full binding curves were generated for the three different species. Except for BBB00497, all VHHs were cross-reactive for mouse, human and cynomolgus MOG. Most cross-reactive VHHs showed a higher affinity towards hMOG compared to mMOG or cMOG (Table 1) (Fig. 2A-C). Two cross-reactive VHHs with a more than 10-fold difference (BBB00498: EC50 3.9 nM, BBB00500: EC50 67.4 nM) in affinity for mMOG were selected for *in vivo* evaluation.

**Figure 2.**
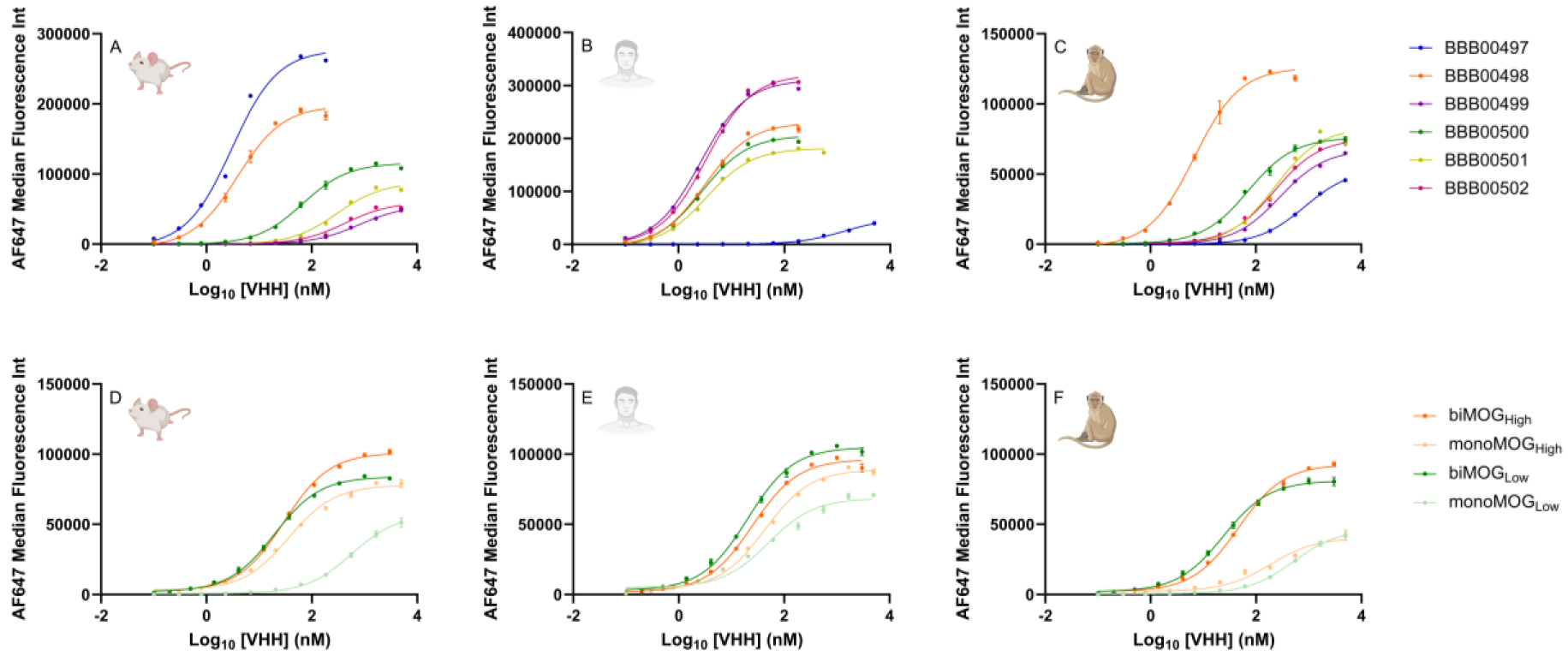
mMOG/hMOG/cMOG affinities of VHHs and antibodies. Flow cytometry binding curves of VHHs and antibodies to CHO cells overexpressing mouse MOG **(A/D)**, human MOG **(B/E)**, and cynomolgus MOG **(C/F)**. Data represents mean ± SEM (BBB00498-BBB00500: n = 2, BBB00497-BBB00499-BBB00500-BBB00501-BBB00502: n = 1, biMOG_High_: n = 2; monoMOG_High_, biMOG_Low_, monoMOG_Low_: n = 3).

**Table 1.** mMOG/hMOG/cMOG affinities of VHHs and antibodies. EC50 values (nM) of flow cytometry on CHO cells overexpressing mouse MOG, human MOG, and cynomolgus MOG.

| VHH/Ab | Mouse EC<sub>50</sub> (nM) | Human EC<sub>50</sub> (nM) | Cynomolgus EC<sub>50</sub> (nM) |
| --- | --- | --- | --- |
| BBB00497 | 3.1 | > 1000 | 844.5 |
| BBB00498 | 3.9 | 3.1 | 6.8 |
| BBB00499 | 800.6 | 2.6 | 285.5 |
| BBB00500 | 67.4 | 2.8 | 71.9 |
| BBB00501 | 304.1 | 3.4 | 234.2 |
| BBB00502 | 434.7 | 3.3 | 225.3 |
| biMOG<sub>High</sub> | 29.6 | 25.0 | 46.33 |
| monoMOG<sub>High</sub> | 36.5 | 43.3 | 184.0 |
| biMOG<sub>Low</sub> | 19.6 | 20.3 | 24 |
| monoMOG<sub>Low</sub> | 573.8 | 50.28 | 519.6 |

### Antibody engineering and in vitro binding confirmation

In a previous study, we showed that Nb62, an in-house discovered VHH binding to mTfR, was able to deliver a therapeutic anti-BACE1 antibody (1A11) to the CNS and elicit a short-term therapeutic effect by decreasing Aβ_1-40_ levels in the brain (17). BACE1 is an enzyme involved in the cleavage of amyloid precursor protein and the formation of Aβ peptides, presenting a therapeutic readout for brain penetration of antibody constructs. To prolong the therapeutic effect, this 1A11-Nb62 antibody-VHH fusion is supplemented with one or two anti-MOG VHHs with different affinities (High: BBB00498, Low: BBB00500) (Fig. 1). Binding of these antibody-VHH fusions to m/h/cMOG was evaluated using flow cytometry (Fig. 2D-F). Three out of four constructs show a similar affinity ranging from 19.6 to 36.5 nM (monoMOG_High_:monoTfR:BACE1, biMOG_Low_:monoTfR:BACE1 and biMOG_High_:monoTfR:BACE1), while one of the constructs had a 15-30 fold lower affinity (monoMOG_Low_:monoTfR:BACE1; 573.8 nM) (Fig. 2D) (Table 1). The affinity of similar antibody constructs for mTfR was determined to be 37 nM in a previous study by biolayer interferometry (17).

### MOG binding valency and affinity affects in vivo PK/PD

To investigate the effect of anti-MOG VHH affinity and avidity on CNS half-life, 5 different antibodies (monoTfR:BACE1, monoMOG_Low_:monoTfR:BACE1, monoMOG_High_:monoTfR:BACE1, biMOG_Low_:monoTfR:BACE1, biMOG_High_:monoTfR:BACE1), were tested *in vivo* at a dose of 80 nmol/kg. Plasma and brain samples were collected from treated mice one and three days post treatment. Antibody concentrations in plasma were similar for all antibodies and decreased rapidly, as expected given the high TfR expression in multiple peripheral organs, resulting in a peripheral sink for these antibodies (Fig. 3A). Aβ_1-40_ levels in plasma decreased significantly for all antibodies compared to PBS treated control mice at both evaluated timepoints (Fig. 3B). Brain PK was similar for all antibodies one day post treatment. However, three days post treatment, antibody concentrations in brain were significantly higher for all MOG binding antibodies except monoMOG_Low_, compared to the monoTfR:BACE1 control (biMOG_High_: p < 0.0001, monoMOG_High_: p = 0.0023, biMOG_Low_: p < 0.0001, monoMOG_Low_: p = 0.1689). biMOG_Low_:monoTfR:BACE1 showed the greatest improvement in brain PK (Fig. 3C). Aβ_1-40_ levels in brain decreased around 40 % compared to PBS treated control mice for all antibodies one day post treatment and for all MOG binding antibodies three days post treatment. The therapeutic effect diminished already three days post treatment for the monoTfR:BACE1 control (Fig. 3D). 40 % decrease in brain Aβ_1-40_ levels is assumed to be the maximum effect possible with antibody 1A11, as the same decrease is observed after direct intracranial injection. Despite the PK differences between the different MOG binding constructs, no significant difference is observed in the therapeutic effect. This could probably be explained by a saturation of the PD effect with all brain concentrations achieved except for the noMOG control.

**Figure 3.**
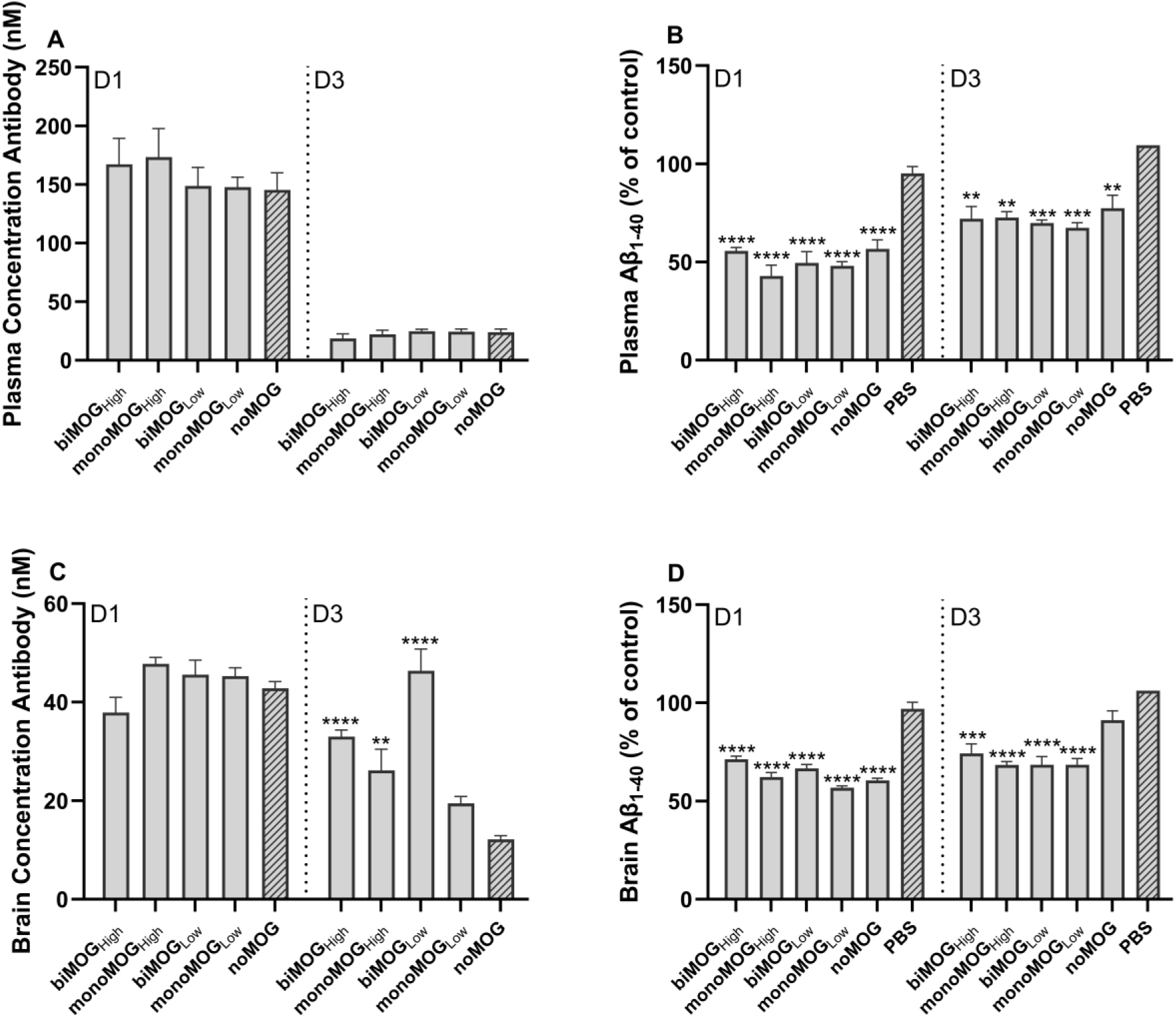
MOG binding valency and affinity affects in vivo PK/PD. Mice received a single intravenous injection (80nmol/kg) with PBS (control), biMOG_High_:monoTfR:BACE1, monoMOG_High_:monoTfR:BACE1, biMOG_Low_:monoTfR:BACE1, monoMOG_Low_:monoTfR:BACE1 or monoTfR:BACE1 (= noMOG). PK/PD were evaluated one and three days post treatment. **(A)** Plasma concentration of treatment antibodies, fast plasma clearance was observed for all antibodies. **(B)** Effect of treatment on Aβ_1-40_ levels in plasma, a similar decrease in plasma Aβ_1-40_ levels was observed for all antibodies. Three days post treatment the decrease was less pronounced then one day post treatment. **(C)** Brain concentration of treatment antibodies, no major differences were observed one day post treatment while MOG binding significantly increased brain antibody levels three days post treatment. biMOG_Low_ showed the highest antibody levels in brain. **(D)** Effect of treatment on Aβ_1-40_ levels in brain, similar decrease in Aβ_1-40_ levels was observed one and three days post treatment for all MOG binding antibodies. Bar graphs represent mean ± SEM (n = 4 per group). Statistical test: two-way ANOVA with Dunnett’s multiple comparisons test compared to monoTfR:BACE1 injected mice for antibody concentrations and to PBS injected control mice for Aβ_1-40_ levels. (*p<0.05, **p<0.01, ***p<0.001, ****p<0.0001)

### Enhanced and prolonged brain uptake in mice of biMOGLow:monoTfR:BACE1

As biMOG_Low_:monoTfR:BACE1 was the best performing antibody construct in terms of brain PK, this antibody was selected for further *in vivo* evaluation. biMOG_Low_:monoTfR:BACE1 was compared head-to-head with monoTfR:BACE1 at a dose of 80 nmol/kg and 160 nmol/kg, and a follow-up period of maximum 14 days. Plasma PK and PD was similar for both antibodies and characterized by a fast clearance, 7 days post treatment no antibody or decrease in Aβ_1-40_ levels could be detected in plasma anymore (Fig. 4B and 4C). However, binding to MOG leads to a significantly increased brain concentration over the whole two-week time period (p < 0.0001). Even with a lower dose of biMOG_Low_:monoTfR:BACE1, significantly higher brain concentrations were achieved than with a high dose of monoTfR:BACE1 (p = 0.0001) (Fig. 4D). This is translated in an increased therapeutic effect, i.e. significantly lower Aβ_1-40_ levels, up to one week post treatment (high dose: p = 0.0208, low dose: p = 0.0413) (Fig. 4E). The increased antibody concentration when binding to MOG was also observed in spinal cord (Fig. S1B).

**Figure 4.**
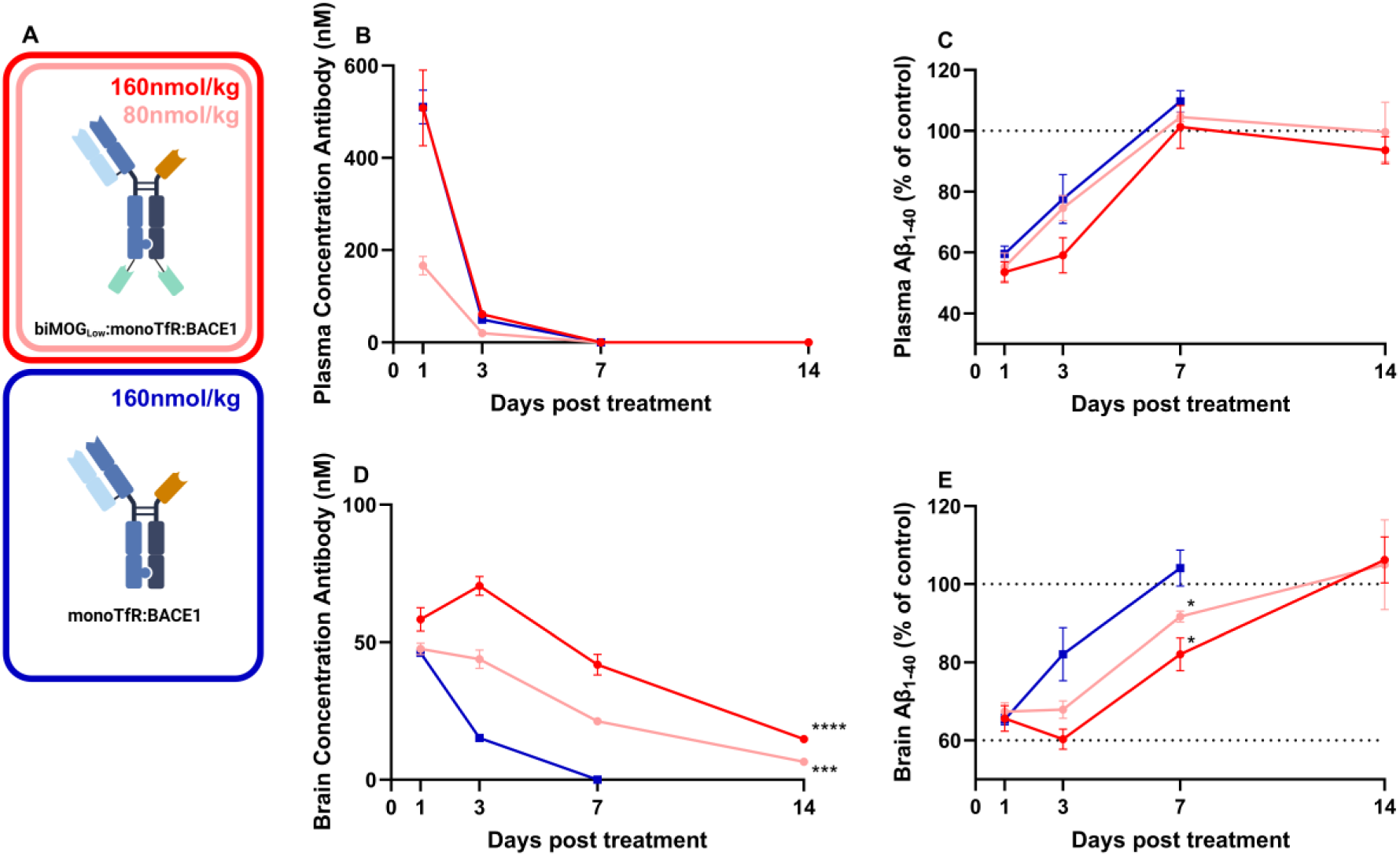
Enhanced and prolonged brain uptake in mice of biMOG_Low_:monoTfR:BACE1. Mice received a single intravenous injection of monoTfR:BACE1 (blue: 160 nmol/kg) or biMOG_Low_:monoTfR:BACE1 (pink: 80 nmol/kg, red: 160 nmol/kg) or PBS (control). PK/PD were evaluated over a period of 7-14 days. **(A)** Schematic representation of biMOG_Low_:monoTfR:BACE1 and monoTfR:BACE1 . **(B)** Plasma concentration of treatment antibodies over time showing fast peripheral clearance. **(C)** Effect of treatment on Aβ_1-40_ levels in plasma, a decrease in therapeutic effect is observed over time. **(D)** Brain concentration of treatment antibodies indicating a higher brain antibody exposure with biMOG_Low_:monoTfR:BACE1. **(E)** Effect of treatment on Aβ1-40 levels in brain, a prolonged therapeutic effect is observed with biMOG_Low_:monoTfR:BACE1 compared to monoTfR/BACE1. Curves represent mean ± SEM (n = 3-8 per group). Statistical test: Two-way ANOVA, all conditions compared to monoTfR:BACE1. **(D)** Significant treatment*time interaction effect for high dose (Red, p < 0.0001) and low dose (Pink, p = 0.0001). **(E)** Significant treatment*time interaction effect for high dose (Red, p = 0.0208) and low dose (Pink, p = 0.0413), only points considered till day 7. (*p<0.05, **p<0.01, ***p<0.001, ****p<0.0001)

### Active shuttling required to reach therapeutically relevant brain concentrations

To avoid the peripheral sink of antibody mediated by TfR binding, MOG binding antibody-based constructs without active shuttling mechanism were generated to evaluate if brain accumulation of antibodies after passive diffusion is sufficient to elicit a therapeutic effect. Peripheral clearance of biMOG_Low_:monoGFP:BACE1 and monoGFP:BACE1 was reduced compared to the TfR binding counterparts. Plasma concentrations one day post treatment were four fold higher and antibodies were still detectable in plasma after 14 days (Fig. 5B). Consistently, Aβ_1-40_ levels in plasma were decreased at all evaluated timepoints (Fig. 5C). Even without active shuttling there is a significant difference between MOG binding and no MOG binding antibodies (p = 0.0001). 14 days post treatment the levels of biMOG_Low_:monoGFP:BACE1 were 5-fold higher then monoGFP:BACE1 (Fig. 5D). However, brain concentrations were too low to elicit any therapeutic effect (Fig. 5E). A similar antibody PK profile for these constructs was observed in spinal cord (Fig. S1C).

**Figure 5.**
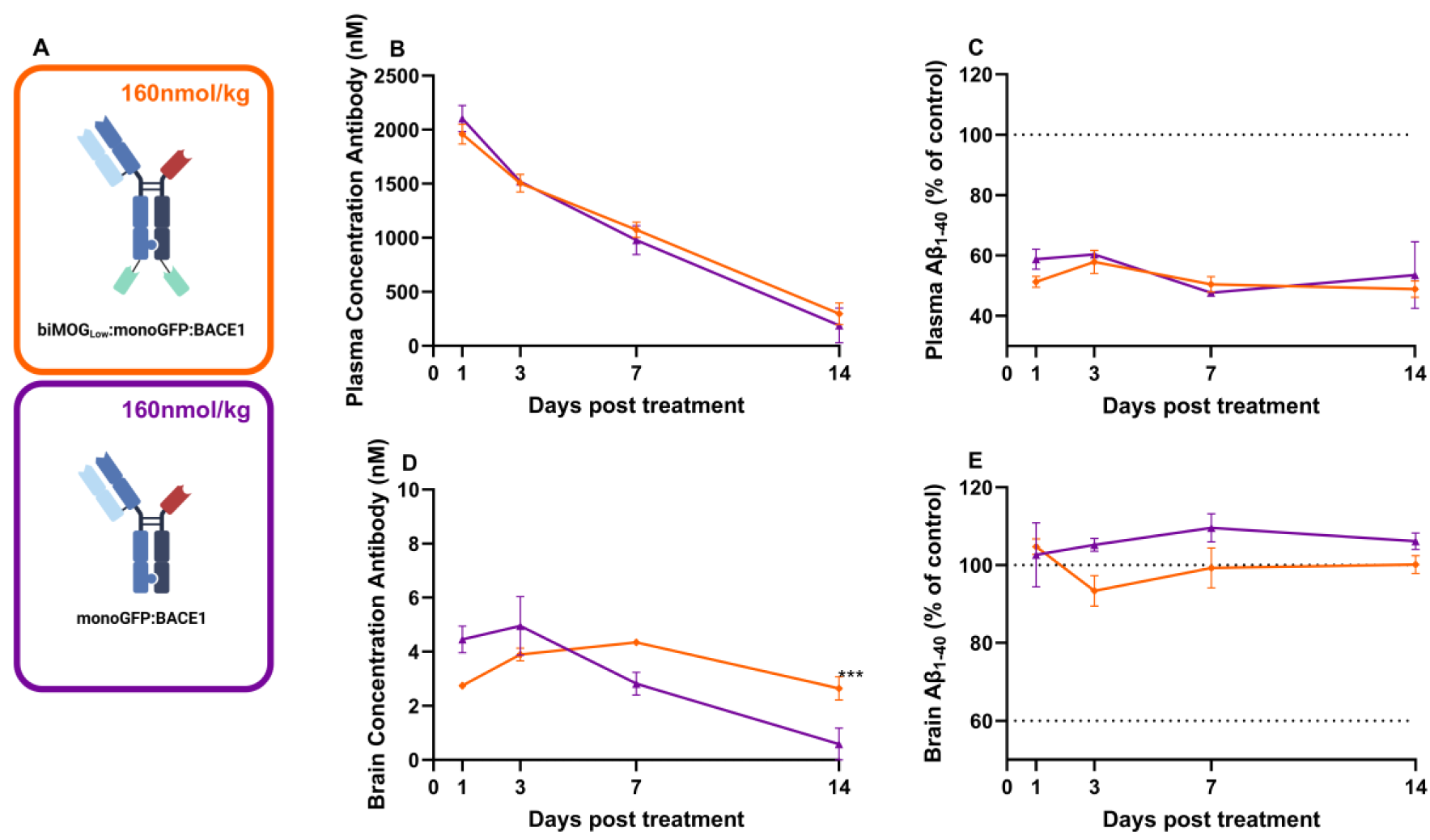
Brain and plasma PK/PD of MOG binding without active shuttling. Mice received a single intravenous injection of 160 nmol/kg of monoGFP:BACE1 (purple) or biMOG_Low_:monoGFP:BACE1 (orange) or PBS (control). PK/PD were evaluated over a period of 14 days. **(A)** Schematic representation of biMOG_Low_:monoGFP:BACE1 and monoTfR:BACE1 . **(B)** Plasma concentration of treatment antibodies over time showing a slower peripheral clearance compared to TfR binding equivalents. **(C)** Effect of treatment on Aβ_1-40_ levels in plasma, the decrease in plasma Aβ_1-40_ levels remained constant over time. **(D)** Brain concentration of treatment antibodies indicating a longer half-life for biMOG_Low_:monoGFP:BACE1 but overall very low brain concentrations. **(E)** Effect of treatment on Aβ_1-40_ levels in brain, no therapeutic effect is observed. Curves represent mean ± SEM (n = 3-6 per group). Statistical test: Two-way ANOVA. **(D)** Significant treatment*time interaction effect (p = 0.0001). (*p<0.05, **p<0.01, ***p<0.001, ****p<0.0001)

### Brain antibody kinetics can be modulated by Fab choice

Above described results were all based on therapeutic antibodies binding BACE1. However, the target of the therapeutic Fab-arm could potentially influence antibody kinetics, for instance by target mediated clearance. To assess the contribution of BACE1 mediated clearance, the BACE1 binding Fab arm was exchanged for a Fab fragment binding to SARS-CoV-2 spike protein (SarsCov), a target not present in the animal model used. Plasma PK remained unchanged compared to their BACE1 binding equivalents (Fig. 6B). While there was almost no difference in brain PK of monoTfR:BACE1 vs monoTfR:SarsCov, a major improvement was observed for biMOG_Low_:monoTfR:SarsCov compared to biMOG_Low_:monoTfR:BACE1 (Fig. 6C). Brain concentrations for both constructs peaked 3 days post treatment but were almost 2-fold higher when eliminating the BACE1 binding. Afterwards, brain antibody concentration decreased to approximately 40 nM at day 21 post treatment, after which a minimal decline was observed up until 49 days post treatment, the final timepoint investigated in this study (Fig. 6C). Also, biMOG_Low_:monoGFP:SarsCov showed increased brain exposure reaching concentrations of 10 nM in the brain (Fig. 6D). Similar results were obtained for spinal cord PK (Fig. S1B-C). When comparing brain antibody exposure (area under the curve (AUC)) over all evaluated treatments and timepoints, biMOG_Low_:monoTfR:SarsCov showed a 4-fold higher exposure then biMOG_Low_:monoTfR:BACE1, a 15-fold higher exposure then monoTfR:SarsCov, the current state-of-the-art for brain antibody delivery, and a 63-fold higher exposure then monoGFP:BACE1, a negative control antibody (Fig. S2).

**Figure 6.**
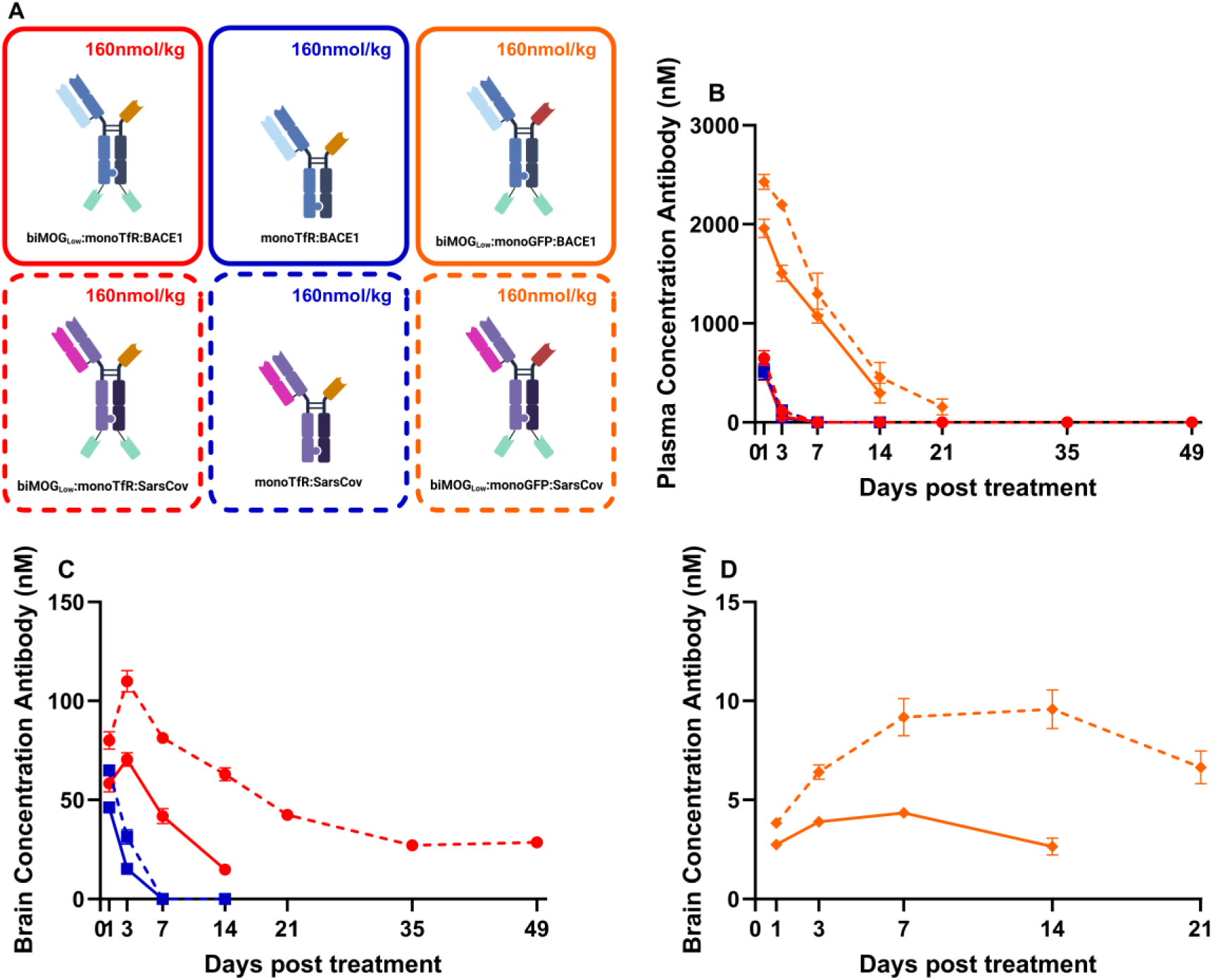
Plasma and brain PK of antibodies with non-targeted Fabs. Mice received a single intravenous injection of 160 nmol/kg of monoTfR:SarsCov (dashed blue), biMOG_Low_:monoTfR:SarsCov (dashed red) or biMOG_Low_:monoGFP:SarsCov (dashed orange). PK/PD were evaluated over a period of 7-14 days. **(A)** Schematic representation of the antibodies . **(B)** Plasma concentration of treatment antibodies over time showing similar profiles for SarsCov and BACE1 binding antibodies. **(C)** Brain concentration of actively shuttled treatment antibodies indicating a higher brain antibody exposure with biMOG_Low_:monoTfR:SarsCov, which is still detectable in brain 49 days post treatment. **(D)** Brain concentration of antibodies without active shuttling, showing an increased brain exposure with non-targeting Fabs. Curves represent mean ± SEM (n = 3-6 per group).

### biMOGLow:monoTfR and noMOG:monoTfR have distinct brain distribution

Widefield imaging of brain sections from mice treated with a single 160 nmol/kg antibody dose 3 days post treatment, immunostained for human IgG (hIgG) revealed insights into the brain biodistribution of biMOG_Low_:monoTfR:SarsCov/BACE1 vs monoTfR:SarsCov/BACE1. Low magnification sagittal sections showed broad distribution of monoTfR:SarsCov, while biMOG_Low_:monoTfR:SarsCov was more distributed to white matter (Fig. 7A). Co-staining for MOG indicated clear overlap between MOG containing regions and biMOG_Low_:monoTfR:SarsCov (Fig. 7A). In general, brain tissue from mice administered monoTfR:SarsCov exhibited markedly less hIgG immunoreactivity than biMOG_Low_:monoTfR:SarsCov, consistent with the lower concentration of this antibody in brain three days post treatment. Magnifications of a brain region from figure 7A visualizing part of the hippocampus, striatum, corpus callosum and cortex, demonstrate a more vascular staining pattern of monoTfR:SarsCov compared to biMOG_Low_:monoTfR:SarsCov, indicated by co-localization with lectin staining (Fig. 7B). Similar observations were found for monoTfR:BACE1 and biMOG_Low_:monoTfR:BACE1 (Fig. S3A-B).

**Figure 7.**
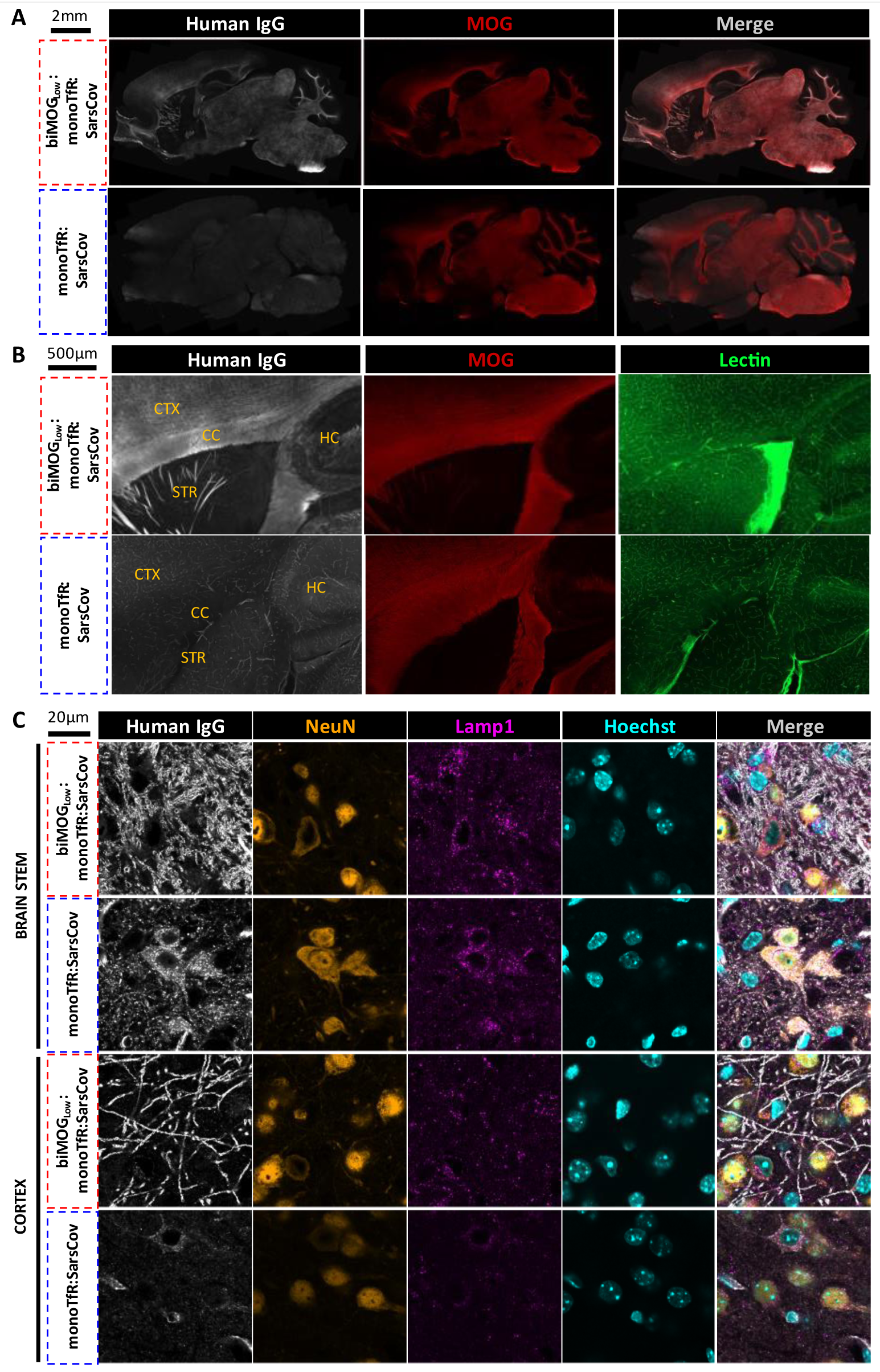
Human IgG localization in mouse brains. Immunohistochemistry of mouse brain sagittal sections 3 days post single 160 nmol/kg IV dose of biMOG_Low_:monoTfR:SarsCov or monoTfR:SarsCov. Representative images are shown from n = 3 animals/group, n = 2 IHC sections/animal. **(A)** Low magnification imaging of mouse brain sagittal sections immunostained for human IgG (white) and MOG (red) showing distinct distribution profiles. **(B)** Zoom of overview scans in A, displaying part of the cortex (CTX), corpus callosum (CC), hippocampus (HC) and striatum (STR) stained for human IgG (white), MOG (red) and lectin (green), demonstrate a more vascular staining pattern of monoTfR:SarsCov compared to biMOG_Low_:monoTfR:SarsCov. **(C)** Confocal imaging of cortex and brain stem, immunostained for human IgG (white), NeuN (orange), Lamp1 (Magenta) and Hoechst (Cyan) indicating more neuronal internalization of monoTfR:SarsCov compared to biMOG_Low_:monoTfR:SarsCov.

Subcellular localization of hIgG in these brains was further examined using super-resolution confocal imaging in cortex and brain stem, a region low and high in MOG expression respectively. Sections were co-stained with the neuronal marker NeuN and a lysosomal marker Lamp1. Prominent cellular internalization in neurons and clear colocalization with neuronal lysosomes was observed for monoTfR:SarsCov in both brain regions while this was absent for biMOG_Low_:monoTfR:SarsCov. The latter showed clear distribution to myelinated fibers in both regions (Fig. 7C). BACE1 binding did not alter these findings (Fig. S3C).

### monoTfR:SarsCov and biMOGLow:monoTfR:SarsCov exhibit similar peripheral biodistribution profiles

To investigate the effect of MOG binding on whole-body biodistribution, antibody constructs were radiolabeled with the PET isotope zirconium-89. Non-targeting Fabs were used to minimize confounding effects on biodistribution. As such, antibody constructs biMOG_Low_:monoTfR:SarsCov and monoTfR:SarsCov were derivatized with the chelator DFO* via random lysine coupling. The radiolabeling was performed under mild conditions, achieving a radiochemical purity of > 99%, as verified by radio-SE-HPLC. The stability of these radio-immunoconjugates was confirmed *in vitro* via iTLC, with the MOG-binding construct showing 89% stability in formulation buffer at 4°C and 86% in normal mouse serum at 37°C after 7 days, while the non-MOG-binding construct demonstrated 91% stability in formulation buffer at 4°C and 84% in normal mouse serum at 37°C over the same period.

Healthy mice were injected with a single dose of either [^89^Zr]Zr-DFO*-biMOG_Low_:monoTfR:SarsCov or [^89^Zr]Zr-DFO*-monoTfR:SarsCov to assess the impact of MOG binding on biodistribution. Whole-body PET imaging was performed on live animals, while *ex vivo* biodistribution studies were conducted on perfused animals at various time points post treatment. The *in vivo* PET imaging data (Fig. 8B) were consistent with the *ex vivo* biodistribution results (Fig. 8A), showing no remarkable differences in peripheral uptake between the two constructs (Fig. S4). In both groups, peripheral distribution was primarily localized to the liver, spleen, kidneys, and bone.

**Figure 8.**
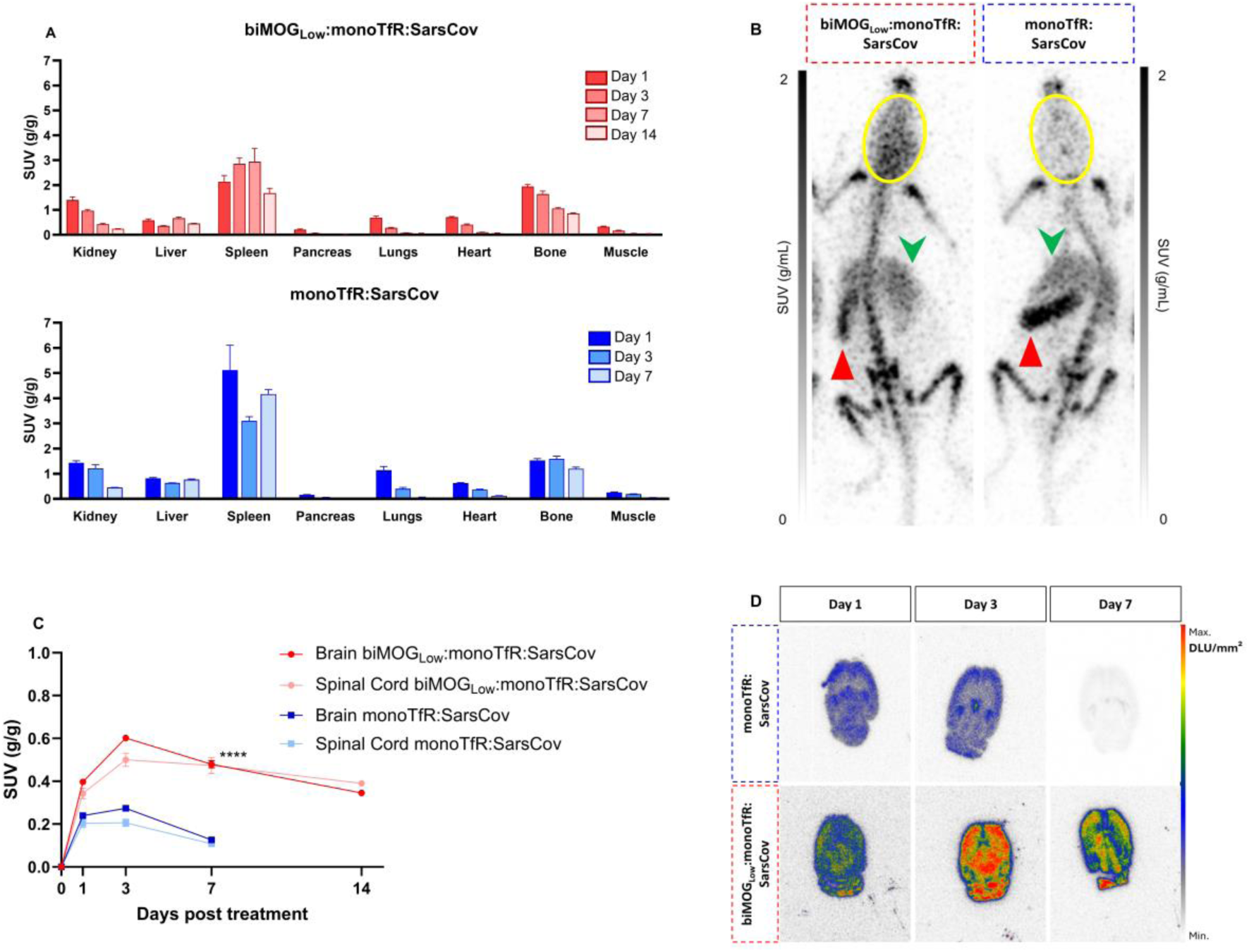
Central and peripheral biodistribution of radiolabeled antibodies. Mice received a single dose of [^89^Zr]Zr-DFO*-biMOG_Low_:monoTfR:SarsCov or [^89^Zr]Zr-DFO*-monoTfR:SarsCov. **(A)** Ex vivo biodistribution, following transcardial perfusion, radiolabeled antibodies were detected in different organs by γ-counting. Bar graphs represent mean standardized uptake value (SUV) (g/g) ± SEM (n = 3-5 per group). **(B)** Representative PET maximum intensity projections of mice (head-first prone position) 7 days post treatment, scaled to SUV (g/mL) for consistent visual comparison. Peripheral uptake is primarily observed in the spleen (red upwards triangle), liver (green downwards arrow), and bone, displaying a similar distribution profile in both groups. However, uptake in the brain region (outlined with a yellow oval) shows increased brain antibody levels in the mouse receiving the biMOG_Low_:monoTfR:SarsCov. **(C)** CNS biodistribution, following transcardial perfusion, radiolabeled antibodies were detected in different organs by γ-counting. Brain and spinal cord levels of radiolabeled antibodies, indicating a higher central antibody exposure with biMOG_Low_:monoTfR:SarsCov. Curves represent mean ± SEM (n = 3-5 per group). Statistical test: Two-way ANOVA, significant treatment*time interaction effect between different treatments in both brain and spinal cord, only points considered till day 7. (**** p<0.0001) **(D)** Representative autoradiography scans of brain slices, showing a higher brain uptake for biMOG_Low_:monoTfR:SarsCov compared to monoTfR:SarsCov.

While peripheral biodistribution profiles were similar between biMOG_Low_:monoTfR:SarsCov and monoTfR:SarsCov (control), there was a notable difference in CNS uptake. For [^89^Zr]Zr-DFO*-monoTfR:SarsCov , brain uptake peaked at 3 days post treatment with an SUV of 0.27 ± 0.001 which declined to 0.13 ± 0.002 at day 7. In contrast, [^89^Zr]Zr-DFO*-biMOG_Low_:monoTfR:SarsCov, also peaked at 3 days post treatment but with a higher SUV of 0.60 ± 0.010 and maintained significantly higher tracer retention (p < 0.0001) up to 14 days, with an SUV value of 0.34 ± 0.008 (Fig. 8C). Similar findings were observed for the spinal cord.

Autoradiography of brain slices from perfused animals was conducted to evaluate the regional distribution of the radiolabeled antibody constructs in the brain (Fig. 8D). [^89^Zr]Zr-DFO*-monoTfR:SarsCov exhibited higher binding to cortical brain regions, while [^89^Zr]Zr-DFO*-biMOG_Low_:monoTfR:SarsCov showed distribution to multiple brain structures and a higher and more prolonged brain uptake compared to the monoTfR:SarsCov, in line with PET and biodistribution experiments.

These results indicate that incorporating MOG-binding VHHs onto a TfR-binding antibody construct does not alter peripheral biodistribution but substantially increases the CNS exposure of the antibody.

### Brain expression levels of antibody targets

As a first safety precaution, the effect of the antibody constructs on their respective targets was evaluated by measuring the expression levels of the different targets at multiple timepoints post treatment. Because western blot is a semi-quantitative technique and to reduce the probability of a type I error, the statistical significance level was set at 1 %. MOG levels remained stable over time for monoTfR:BACE1, biMOG_Low_:monoTfR:BACE1 and biMOG_Low_:monoTfR:SarsCov (Fig. 9A). Animals receiving biMOG_Low_:monoTfR:BACE1 showed increasing brain levels of BACE1 over time while this was not observed with monoTfR:BACE1 or biMOG_Low_:monoTfR:SarsCov (Fig. 9B). Brain TfR levels were decreased in mice treated with monoTfR:BACE1, in contrast to an increasing trend of brain TfR levels when treated with biMOG_Low_:monoTfR:SarsCov (Fig. 9C).

**Figure 9.**
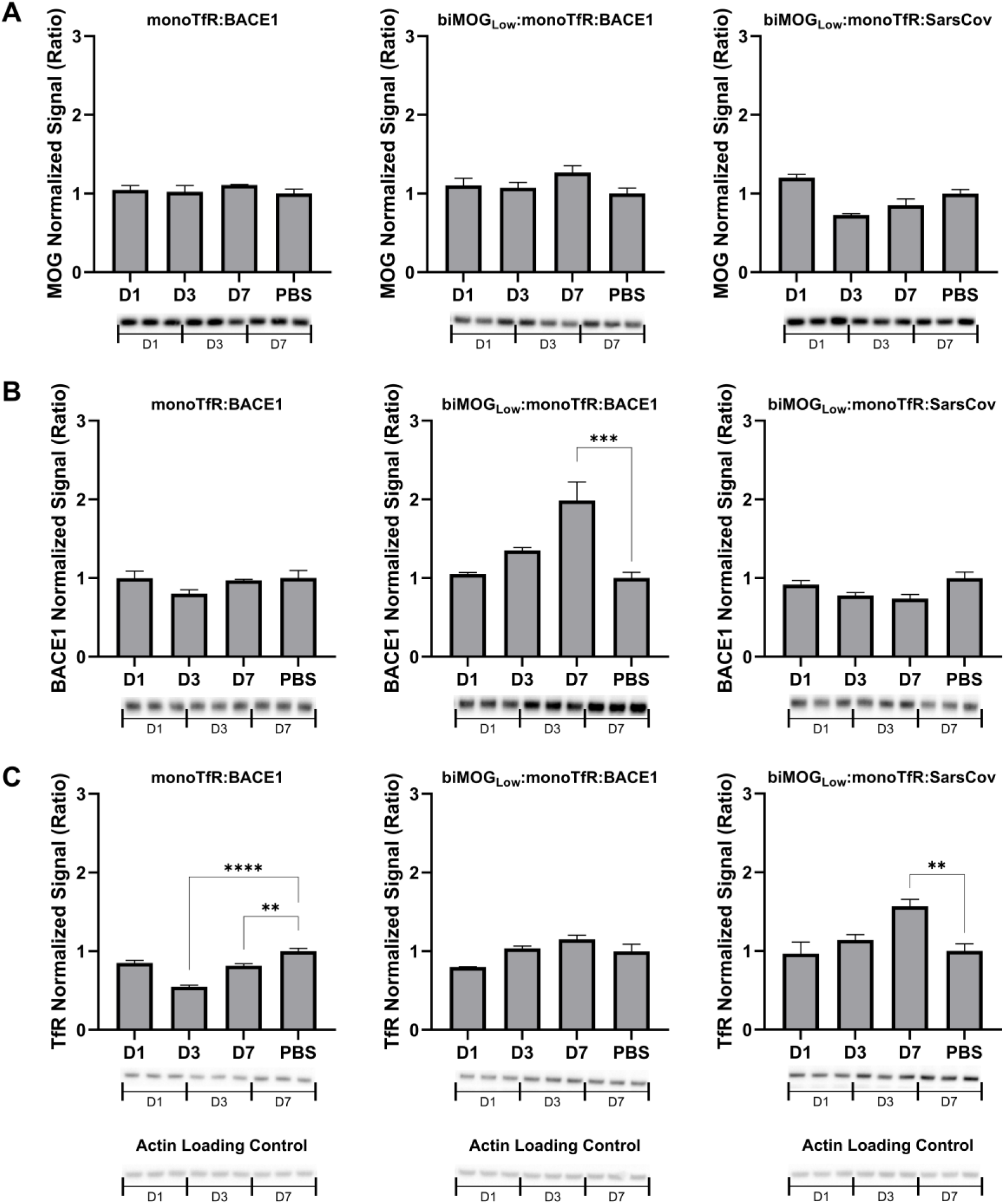
Target protein expression levels in brain homogenates. Western blots and quantification of **(A)** MOG, **(B)** BACE1 and **(C)** TfR in brain homogenates of mice treated with a single IV injection of 160 nmol/kg biMOG_Low_:monoTfR:BACE1, biMOG_Low_:monoTfR:SarsCov or monoTfR:BACE1. Target expression levels are evaluated 1, 3, and 7 days post treatment. Bar graphs represent mean ± SEM (n = 3 per group) of intensity signal normalized to actin loading control, and displayed as a ratio compared to PBS treated control mice. Statistical test: one-way ANOVA with Dunnett’s multiple comparisons test compared to PBS injected control mice, statistical significance level set to 1 %. (**p<0.01, ***p<0.001, ****p<0.0001)

## DISCUSSION

Drug discovery for neurological disorders is hampered by the restricted access of large molecules to the brain due to the presence of the BBB (26). Extensive academic and industrial research focuses on the use of RMT to shuttle therapies into the brain (7,8,10,17,18,27). TfR mediated shuttling is the most-well studied approach and has successfully entered the clinic by the approval of Izcargo® in Japan (12–14). While brain concentrations are significantly increased when using TfR-shuttling, this effect is short-term requiring frequent dosing (17). To overcome this challenge, we supplemented this brain delivery strategy with binding to a brain specific protein, MOG, leading to a drastically improved central half-life of the antibodies.

First, VHHs targeting the brain specific protein MOG were discovered. VHHs originate from heavy-chain only antibodies found in camelids and can easily be fused to a wide variety of therapeutics, including antibodies, allowing the design of multispecific antibodies (28). While species cross-reactivity remains a challenge for certain antibody/VHH targets such as TfR (18), the discovered MOG binding VHHs are cross-reactive for mouse, human and cynomolgus MOG.

The optimal MOG affinity for brain retention is not known. Strong binding might lead to more accumulation, but potentially less free antibody concentration in the brain and hence lower efficacy; vice versa, lower affinity could lead to lower accumulation, and hence lower efficacy. Indeed, constructs with an apparent affinity between 19.6 to 36.5 nM showed an improved brain accumulation, while an apparent affinity of 573.8 nM seems to be too weak (Fig. 3C). The effect on the brain PD could not be assessed in this study, as a similar maximal inhibition was observed for all four constructs (Fig. 3D). To assess the optimal affinity to reach the optimal PD effects, the *in vivo* experiments should be repeated at a lower dose to potentially see an effect.

A full 14-day PK/PD evaluation was performed for biMOG_Low_:monoTfR:BACE1 and compared to monoTfR:BACE1. While plasma clearance is similar for both antibodies, brain PK significantly improved by binding to MOG. While the non-MOG binding antibody is undetectable in plasma after 7 days, we could still detect MOG binding antibody in the brain 14 days post treatment. Also, the therapeutic effect was prolonged in time, up to 7 days. Interestingly, a discrepancy is observed between brain antibody concentration and therapeutic effect. After 7 days, the brain concentration of biMOG_Low_:monoTfR:BACE1 is the same as the concentration of monoTfR:BACE1 after 1 day. The therapeutic effect on the other hand is lower, ∼20 % decrease in brain Aβ_1-40_ levels at day 7 compared to ∼40 % at day 1, respectively (Fig. 4D/E). This could potentially be explained by an upregulation of BACE1 in mice treated with biMOG_Low_:monoTfR:BACE1 as a coping mechanism for prolonged BACE1 inhibition. Others have confirmed that BACE1 inhibitors can extend the half-life of BACE1 in rat primary cortical neurons (29) and we also observe an increase in BACE1 levels in mouse brain homogenates upon prolonged BACE1 inhibition (Fig. 9B).

By eliminating TfR binding, the peripheral half-life of the antibodies is improved but the concentration reaching the brain is too low to elicit a therapeutic effect. Up to now, the short peripheral half-life of TfR mediated BBB crossing antibodies was mostly considered a disadvantage as the brain PK of such antibodies is following the peripheral PK, leading to fast clearance from the brain. Interestingly, in our approach, the antibody brain PK seems to be more disconnected from the peripheral PK as the antibody remains detectable in the brain when it is already completely cleared in plasma. This scenario of short peripheral half-life combined with long central half-life can be a great advantage for drugs exerting peripheral side effects. These would be short-term due to the fast peripheral clearance, while central, desired therapeutic effects would last longer. Even more, when exchanging the BACE1 binding Fab for a non-targeting Fab (anti-SarsCov), kinetics of brain exposure improve further with an approximately two-fold higher C_max_ and brain antibody levels around 30 nM 49 days post treatment (Fig. 6C). This is a huge advancement for the field as it is the first time that antibodies are detected in brain 7 weeks after a single antibody injection, even at therapeutically relevant concentrations.

Next, we validated the extended brain half-life of MOG binding multispecific antibodies by radio-imaging studies and in parallel evaluated their peripheral biodistribution. This study with ^89^Zr-labeled antibodies confirms the increased uptake and prolonged biological half-life of biMOG_Low_:monoTfR:SarsCov in the CNS. Peripheral biodistribution is similar for both monoTfR:SarsCov and biMOG_Low_:monoTfR:SarsCov, with spleen, bone, kidney and liver being the main organs where zirconium-89 was detected. Spleen and bone marrow uptake can be attributed to the high expression of TfR in these organs (30,31), to antibody metabolism via the reticuloendothelial system (32), or potentially a combination of both. Accumulation in kidney and liver could be related to their function as catabolic organs (33). The increased bone signal can potentially be related to the accumulation of uncomplexed ^89^Zr, known to be a bone seeking agent (34). Indeed, intracellular antibody catabolism occurs in acidic lysosomes, and it is known that zirconium will be released from its chelator under such conditions. Overall, the peripheral biodistribution of both antibodies displays a general profile expected from a ^89^Zr-labeled / TfR targeting antibody construct (35), and no overt differences are observed in peripheral biodistribution between MOG binding and non-MOG binding multispecific antibodies.

When looking more in detail to the central biodistribution, we show that biMOG_Low_:monoTfR:SarsCov displays baseline distribution to white matter, in contrast to the more uniform distribution of monoTfR:SarsCov. The latter shows neuronal internalization and colocalization with lysosomes which is absent for biMOG_Low_:monoTfR:SarsCov, probably contributing to its lower turnover rate and higher antibody concentrations in the brain. The addition of BACE1 binding does not change this localization. However, others have shown that Fab target binding can redirect antibodies to other cell types (9), making it probably more dependent on the affinity balance between the different targets. For monoTfR:SarsCov vascular localization is observed. As this is not the case for biMOG_Low_:monoTfR:SarsCov it cannot be attributed to insufficient brain perfusion. A potential explanation is that part of TfR binding antibodies that reached the brain remains attached to the brain vasculature at the brain parenchymal side, while MOG binding lowers the free concentration in the brain, resulting in an enhanced release from the vasculature.

In addition, our immunofluorescence studies show an altered brain and cellular distribution of TfR-mediated BBB crossing antibodies when fused to anti-MOG VHHs. The observed intracellular (without anti-MOG VHHs) to extracellular relocation (with anti-MOG VHHs) of TfR-mediated BBB crossing antibodies might be beneficial to increase drug concentration around extracellular targets but might be less preferred for treatment paradigms where internalization is required (e.g. for antisense oligonucleotide delivery or enzyme replacement therapies for lysosomal storage diseases). Finally, MOG targeting might be additionally beneficial for treatment of pathologies affecting myelination like e.g. multiple sclerosis, as also here the drug might be more relocated towards the location to be treated. Importantly, whether or not additional MOG binding is beneficial for a given treatment will need to be evaluated on a case by case basis and fundamental understanding of disease mechanisms and treatment will be key to determine the optimal brain delivery approach.

To induce experimental allergic encephalomyelitis in mouse models, researchers immunize mice with recombinant MOG or peptides thereof (36). Although it has been shown that the auto-immune response is mainly CD4+ T cell related, also anti-MOG antibodies contribute to the disease (37). Therefore, an important consideration is the safety of multispecific antibodies also targeting MOG. For this reason, the antibodies used in this study contain effector and complement knock-out mutations in the Fc region of the antibodies. In a preliminary study, we have observed that mice treated with effector and complement competent version of these antibodies showed an approximately 20 % decrease in brain size 3 days post treatment (data not shown). Given this limitation, the current brain half-life extension approach is probably not suitable for treatments with antibodies relying on immune activation by engagement of the Fcγ receptor or binding to the C1q complement component for their therapeutic effect (38).

In this study, we conducted a limited toxicity test, by evaluating target expression levels in brain homogenates. There is no effect on the brain levels of MOG, however TfR is downregulated after treatment with monoTfR:BACE1 and upregulated with biMOG_Low_:monoTfR:SarsCov. Decreased brain TfR levels after treatment with antibodies with a high affinity for TfR were previously reported and attributed to constitutive degradation of TfR in lysosomes (39,40). However, adding MOG binding seems to inverse this effect. Lysosomal degradation is probably reduced due to the strong affinity for MOG upon entering the brain. Although TfR levels are higher than baseline in mice treated with biMOG_Low_:monoTfR:SarsCov, this increasing trend remains so far unexplained. Furthermore, BACE1 levels are upregulated in mice treated with biMOG_Low_:monoTfR:BACE1. As this is not observed with biMOG_Low_:monoTfR:SarsCov, it is hypothesized that this is a compensation mechanism for continuous inhibition of BACE1.

In order to allow further clinical development of this anti-MOG VHH, it needs to be sequence optimized and ideally the affinity gap between human and mouse needs to be reduced. In this respect, we have humanized the current lead VHH and we removed two potential post translational modification sites in CDR2 and CDR3 respectively. The resulting lead VHH is highly stable (melting temperature > 90 °C) and the affinity gap is reduced from 25-fold to 4-fold by reducing the affinity for human MOG while keeping the affinity for mouse MOG unchanged (data not shown). Moreover, the experiments with radiolabeled antibodies align with other pharmacokinetic studies that measured antibody concentrations in brain homogenates using ELISA. This consistency further underscores the robustness of the findings and validates a non-invasive approach to evaluate the PK behavior of these shuttling antibody constructs through in-vivo PET/CT imaging.

## CONCLUSION

This study reported the development of mouse/human/cyno cross-reactive anti-MOG VHHs and their ability to drastically increase brain exposure of antibodies, resulting in both a higher C_max_ and a longer brain half-life. Combining MOG and TfR binding leads to distinct PK, biodistribution, and brain exposure, differentiating it from the highly investigated TfR-shuttling. It is the first time such long brain antibody exposure is demonstrated after one single dose. This new approach of adding a binding moiety for brain specific targets to RMT shuttling antibodies is a huge advancement for the field and paves the way for further research into brain half-life extension, exploiting other brain specific targets, or the combination with other active shuttling mechanisms. These findings represent a major breakthrough in enhancing brain accumulation of biological agents, overcoming a significant hurdle in the development of new treatments for neurological disorders.

## LIST OF ABBREVIATIONS

AUC: area under the curve
BACE1: β-secretase 1
BBB: blood-brain barrier
CC: corpus callosum
CHO: Chinese hamster ovary
cMOG: cynomolgus MOG
CNS: central nervous system
CT: computed tomography
CTX: cortex
DMSO: dimethylsulfoxide
FBS: fetal bovine serum
GFP: green fluorescent protein
HC: hippocampus
hIgG: human IgG
hMOG: human MOG
iTLC: instant thin layer chromatography
mAb: monoclonal antibody
mMOG: mouse MOG
MOG: myelin oligodendrocyte glycoprotein
ON: overnight
PD: pharmacodynamics
PET: positron emission tomography
PFA: paraformaldehyde
PK: pharmacokinetics
R&D: research & development
RMT: receptor-mediated transcytosis
RT: room temperature
STR: striatum
SUV: standardized uptake value
TfR: transferrin receptor
VHH: variable heavy domain of heavy chain

## DECLARATIONS

### Author Contributions

Conceptualization: M.C., T.J., B.D.S. and M.D. Methodology: M.C., T.J., G.B., F.C. and M.D. Investigation: M.C., T.J., J.C., S.L. and C.C. Data curation: M.C. Visualization: M.C. Supervision: M.D. Writing-original draft: M.C. Writing-review and editing: M.C., T.J., J.C., S.L., C.C., G.B., F.C., B.D.S., N.G. and M.D. Funding acquisition: M.C., N.G. and M.D.

## Acknowledgments

We would like to thank the VIB Nanobody Core for performing the immunizations and generating the VHH-displaying phage library. Special thanks to Isabelle Degors and Laura Rué, for their assistance with immunization DNA and phage preparations, and cell line generation, to Danté Van Humbeeck for radiolabeling of antibodies, and to Julie Cornelis for her assistance with *in vivo* radiopharmaceutical studies.

## Ethics Approval

All animal experiments were approved by the KU Leuven Animal Ethics Committee (project P091/2022 and P038/2024) following governmental and EU guidelines.

## REFERENCES

1. WHO Team Mental Health, Brain Health and Substance Use. Optimizing brain health across the life course: WHO position paper [Internet]. 2022 Sep [cited 2024 Feb 22]. Available from: https://www.who.int/publications-detail-redirect/9789240054561

2. Gores M, Lutzmayer S. Fostering Success in CNS Innovation [Internet]. IQVIA; 2024 Apr. Available from: https://www.iqvia.com/library/white-papers/apac-perspective-on-fostering-success-in-cns-innovation

3. Gores M, Lutzmayer S. Two Steps Forward, One Step Back: The Long Road to Success in CNS [Internet]. IQVIA; 2023 Mar [cited 2023 Dec 8]. Available from: https://www.iqvia.com/library/white-papers/two-steps-forward-one-step-back-the-long-road-to-success-in-cns

4. Almutairi MMA, Gong C, Xu YG, Chang Y, Shi H. Factors controlling permeability of the blood– brain barrier. Cell Mol Life Sci. 2016 Jan 1;73(1):57–77.

5. Freskgård PO, Urich E. Antibody therapies in CNS diseases. Neuropharmacology. 2017 Jul 1;120:38–55.

6. Pardridge WM. A Historical Review of Brain Drug Delivery. Pharmaceutics. 2022 Jun;14(6):1283.

7. Wouters Y, Jaspers T, De Strooper B, Dewilde M. Identification and in vivo characterization of a brain-penetrating nanobody. Fluids Barriers CNS [Internet]. 2020 Oct 14 [cited 2021 Jan 13];17. Available from: https://www.ncbi.nlm.nih.gov/pmc/articles/PMC7556960/

8. Kariolis MS, Wells RC, Getz JA, Kwan W, Mahon CS, Tong R, et al. Brain delivery of therapeutic proteins using an Fc fragment blood-brain barrier transport vehicle in mice and monkeys. Science Translational Medicine. 2020 May 27;12(545):eaay1359.

9. Chew KS, Wells RC, Moshkforoush A, Chan D, Lechtenberg KJ, Tran HL, et al. CD98hc is a target for brain delivery of biotherapeutics. Nat Commun. 2023 Aug 19;14(1):5053.

10. Yu YJ, Zhang Y, Kenrick M, Hoyte K, Luk W, Lu Y, et al. Boosting brain uptake of a therapeutic antibody by reducing its affinity for a transcytosis target. Sci Transl Med. 2011 May 25;3(84):84ra44.

11. Yu YJ, Atwal JK, Zhang Y, Tong RK, Wildsmith KR, Tan C, et al. Therapeutic bispecific antibodies cross the blood-brain barrier in nonhuman primates. Science Translational Medicine. 2014 Nov 5;6(261):261ra154–261ra154.

12. Giugliani R, Martins AM, So S, Yamamoto T, Yamaoka M, Ikeda T, et al. Iduronate-2-sulfatase fused with anti-hTfR antibody, pabinafusp alfa, for MPS-II: A phase 2 trial in Brazil. Molecular Therapy. 2021 Mar 27;29(7):2378–86.

13. Sonoda H, Takahashi K, Minami K, Hirato T, Yamamoto T, So S, et al. Treatment of Neuronopathic Mucopolysaccharidoses with Blood–Brain Barrier-Crossing Enzymes: Clinical Application of Receptor-Mediated Transcytosis. Pharmaceutics. 2022 Jun;14(6):1240.

14. Yamamoto R, Kawashima S. [Pharmacological property, mechanism of action and clinical study results of Pabinafusp Alfa (Genetical Recombination) (IZCARGO® I.V. Infusion 10 mg) as the therapeutic for Mucopolysaccharidosis type-II (Hunter syndrome)]. Nihon Yakurigaku Zasshi. 2022;157(1):62–75.

15. Cummings J. Anti-Amyloid Monoclonal Antibodies are Transformative Treatments that Redefine Alzheimer’s Disease Therapeutics. Drugs. 2023;83(7):569–76.

16. Sevigny J, Chiao P, Bussière T, Weinreb PH, Williams L, Maier M, et al. The antibody aducanumab reduces Aβ plaques in Alzheimer’s disease. Nature. 2016 Sep;537(7618):50–6.

17. Wouters Y, Jaspers T, Rué L, Serneels L, De Strooper B, Dewilde M. VHHs as tools for therapeutic protein delivery to the central nervous system. Fluids and Barriers of the CNS. 2022 Oct 3;19(1):79.

18. Rué L, Jaspers T, Degors IMS, Noppen S, Schols D, De Strooper B, et al. Novel Human/Non-Human Primate Cross-Reactive Anti-Transferrin Receptor Nanobodies for Brain Delivery of Biologics. Pharmaceutics. 2023 Jun;15(6):1748.

19. Atwal J, Chen Y, Chiu C, Mortensen D, Meilandt W, Liu Y, et al. A Therapeutic Antibody Targeting BACE1 Inhibits Amyloid-β Production in Vivo. Science translational medicine. 2011 May 25;3:84ra43.

20. Nakano R, Takagi-Maeda S, Ito Y, Kishimoto S, Osato T, Noguchi K, et al. A new technology for increasing therapeutic protein levels in the brain over extended periods. PLOS ONE. 2019 Apr 12;14(4):e0214404.

21. Takahashi N, Nakano R, Maeda S, Ito Y. Antibody capable of binding to myelin oligodendrocyte glycoprotein [Internet]. WO2018123979A1, 2018 [cited 2023 Dec 8]. Available from: https://patents.google.com/patent/WO2018123979A1/en

22. Pardon E, Laeremans T, Triest S, Rasmussen SGF, Wohlkönig A, Ruf A, et al. A general protocol for the generation of Nanobodies for structural biology. Nat Protoc. 2014 Mar;9(3):674–93.

23. Zhou L, Chávez-Gutiérrez L, Bockstael K, Sannerud R, Annaert W, May PC, et al. Inhibition of β-Secretase in Vivo via Antibody Binding to Unique Loops (D and F) of BACE1. J Biol Chem. 2011 Mar 11;286(10):8677–87.

24. Imbrechts M, Maes W, Ampofo L, Van den Berghe N, Calcoen B, Van Looveren D, et al. Potent neutralizing anti-SARS-CoV-2 human antibodies cure infection with SARS-CoV-2 variants in hamster model. iScience. 2022 Aug 19;25(8):104705.

25. Nesspor TC, Kinealy K, Mazzanti N, Diem MD, Boye K, Hoffman H, et al. High-Throughput Generation of Bipod (Fab × scFv) Bispecific Antibodies Exploits Differential Chain Expression and Affinity Capture. Sci Rep. 2020 May 5;10(1):7557.

26. Pardridge WM. Blood-Brain Barrier and Delivery of Protein and Gene Therapeutics to Brain. Front Aging Neurosci [Internet]. 2020 Jan 10 [cited 2021 Jan 12];11. Available from: https://www.ncbi.nlm.nih.gov/pmc/articles/PMC6966240/

27. Niewoehner J, Bohrmann B, Collin L, Urich E, Sade H, Maier P, et al. Increased brain penetration and potency of a therapeutic antibody using a monovalent molecular shuttle. Neuron. 2014 Jan 8;81(1):49–60.

28. Harmsen MM, De Haard HJ. Properties, production, and applications of camelid single-domain antibody fragments. Appl Microbiol Biotechnol. 2007;77(1):13–22.

29. Liu L, Lauro BM, Ding L, Rovere M, Wolfe MS, Selkoe DJ. Multiple BACE1 inhibitors abnormally increase the BACE1 protein level in neurons by prolonging its half-life. Alzheimers Dement. 2019 Sep;15(9):1183–94.

30. Marsee DK, Pinkus GS, Yu H. CD71 (transferrin receptor): an effective marker for erythroid precursors in bone marrow biopsy specimens. Am J Clin Pathol. 2010 Sep;134(3):429–35.

31. Sugyo A, Tsuji AB, Sudo H, Nomura F, Satoh H, Koizumi M, et al. Uptake of 111In-labeled fully human monoclonal antibody TSP-A18 reflects transferrin receptor expression in normal organs and tissues of mice. Oncology Reports. 2017 Mar 1;37(3):1529–36.

32. Tabrizi MA, Tseng CML, Roskos LK. Elimination mechanisms of therapeutic monoclonal antibodies. Drug Discovery Today. 2006 Jan 1;11(1):81–8.

33. Eigenmann MJ, Fronton L, Grimm HP, Otteneder MB, Krippendorff BF. Quantification of IgG monoclonal antibody clearance in tissues. mAbs. 2017 Aug;9(6):1007.

34. Deri MA, Zeglis BM, Francesconi LC, Lewis JS. PET imaging with 89Zr: From radiochemistry to the clinic. Nuclear Medicine and Biology. 2013 Jan 1;40(1):3–14.

35. Stergiou N, Wuensche TE, Schreurs M, Mes I, Verlaan M, Kooijman EJM, et al. Application of 89Zr-DFO*-immuno-PET to assess improved target engagement of a bispecific anti-amyloid-ß monoclonal antibody. Eur J Nucl Med Mol Imaging. 2023;50(5):1306–17.

36. Bittner S, Afzali AM, Wiendl H, Meuth SG. Myelin Oligodendrocyte Glycoprotein (MOG35-55) Induced Experimental Autoimmune Encephalomyelitis (EAE) in C57BL/6 Mice. J Vis Exp. 2014 Apr 15;(86):51275.

37. Berer K, Wekerle H, Krishnamoorthy G. B cells in spontaneous autoimmune diseases of the central nervous system. Mol Immunol. 2011 Jun;48(11):1332–7.

38. Gogesch P, Dudek S, van Zandbergen G, Waibler Z, Anzaghe M. The Role of Fc Receptors on the Effectiveness of Therapeutic Monoclonal Antibodies. Int J Mol Sci. 2021 Aug 19;22(16):8947.

39. Bien-Ly N, Yu YJ, Bumbaca D, Elstrott J, Boswell CA, Zhang Y, et al. Transferrin receptor (TfR) trafficking determines brain uptake of TfR antibody affinity variants. J Exp Med. 2014 Feb 10;211(2):233–44.

40. Do TM, Capdevila C, Pradier L, Blanchard V, Lopez-Grancha M, Schussler N, et al. Tetravalent Bispecific Tandem Antibodies Improve Brain Exposure and Efficacy in an Amyloid Transgenic Mouse Model. Mol Ther Methods Clin Dev. 2020 Aug 21;19:58–77.

