## Supplementary for "Increasing brain half-life of antibodies by additional binding to myelin oligodendrocyte glycoprotein, a CNS specific protein"

Marie-Lynn Cuypers *et al.*

**This file includes:**

Tables S1 to S4

Figures S1 to S4

*Table S1. Primary and secondary antibody dilutions for brain immunohistochemistry.*

| **Target** | **Primary Antibody** | **Secondary Antibody** |
|---|---|---|
| Nuclei | Hoechst 33342<br>(Sigma, B2261)<br>*Dilution 1/5000* | / |
| Human IgG | Goat anti-human IgG (H+L)-AF647<br>(Thermo Fisher Scientific, A21445)<br>*Dilution 1/200* | / |
| Blood vessels | Tomato Lectin Dyelight 488<br>(Thermo Fisher Scientific, L32470)<br>*Dilution 1/50* | / |
| NeuN | Rabbit anti-NeuN<br>(Abcam, ab177487)<br>*Dilution 1/2000* | Goat anti-rabbit IgG (H+L)-AF568<br>(Thermo Fisher Scientific, A11011)<br>*Dilution 1/1000* |
| MOG | Rabbit anti-MOG antibody<br>(Abcam, AB233549)<br>*Dilution 1/500* | Goat anti-rabbit IgG (H+L)-AF568<br>(Thermo Fisher Scientific, A11011)<br>*Dilution 1/1000* |
| LAMP1 | Rat anti-Mouse CD107a<br>(BD Biosciences, 553792)<br>*Dilution 1/250* | Goat anti-rat IgG (H+L)-AF488<br>(Thermo Fisher Scientific, A11006)<br>*Dilution 1/500* |

*Table S2. Primary and secondary antibody dilutions for western blotting.*

| **Target** | **Primary Antibody** | **Secondary Antibody** |
|---|---|---|
| MOG | MOG Polyclonal Antibody, Rabbit<br>(Thermo Fisher Scientific, PA595602)<br>*Dilution 1/2000* | Goat anti-rabbit IgG (H+L)-HRP<br>(Bio-Rad, 170-6515)<br>*Dilution 1/3000* |
| BACE1 | BACE1 (D10E5) Rabbit mAb<br>(Cell Signaling, 5606S)<br>*Dilution 1/500* | Goat anti-rabbit IgG (H+L)-HRP<br>(Bio-Rad, 170-6515)<br>*Dilution 1/1000* |
| TfR | TfR Monoclonal Antibody<br>(Thermo Fisher Scientific, 13-6800)<br>*Dilution 1/1000* | Goat anti-mouse HRP<br>(Agilent, P044701)<br>*Dilution 1/5000* |

*Table S3. Overall statistical comparisons.*

| Figure/Segment | Statistical Test | α level | Variables | F | DF | P value |
|---|---|---|---|---|---|---|
| Figure 3A | | | | | | |
| | *Two-way ANOVA* | 0.05 | Treatment | 0.3694 | 4 | 0.8285 |
| | | | Time | 270.5 | 1 | <0.0001 |
| | | | Interaction | 0.6690 | 4 | 0.6185 |
| Figure 3B | | | | | | |
| | *Two-way ANOVA* | 0.05 | Treatment | 12.75 | 5 | <0.0001 |
| | | | Time | 48.95 | 1 | <0.0001 |
| | | | Interaction | 0.6152 | 5 | 0.6890 |
| Figure 3C | | | | | | |
| | *Two-way ANOVA* | 0.05 | Treatment | 12.77 | 4 | <0.0001 |
| | | | Time | 146.0 | 1 | <0.0001 |
| | | | Interaction | 17.99 | 4 | <0.0001 |
| Figure 3D | | | | | | |
| | *Two-way ANOVA* | 0.05 | Treatment | 20.36 | 5 | <0.0001 |
| | | | Time | 27.97 | 1 | <0.0001 |
| | | | Interaction | 6.323 | 5 | 0.0004 |
| Figure 4B (160 nmol/kg biMOG$_{Low}$:monoTfR:BACE1 vs 160 nmol/kg monoTfR:BACE1) | | | | | | |
| | *Two-way ANOVA (only datapoints until day 7 included)* | 0.05 | Treatment | 0.006111 | 1 | 0.9384 |
| | | | Time | 62.27 | 2 | <0.0001 |
| | | | Interaction | 0.01065 | 2 | 0.9894 |
| Figure 4C (160 nmol/kg biMOG$_{Low}$:monoTfR:BACE1 vs 160 nmol/kg monoTfR:BACE1) | | | | | | |
| | *Two-way ANOVA (only datapoints until day 7 included)* | 0.05 | Treatment | 4.524 | 1 | 0.0454 |
| | | | Time | 32.62 | 2 | <0.0001 |
| | | | Interaction | 0.5539 | 2 | 0.5829 |
| Figure 4D (160 nmol/kg biMOG$_{Low}$:monoTfR:BACE1 vs 160 nmol/kg monoTfR:BACE1) | | | | | | |
| | *Two-way ANOVA* | 0.05 | Treatment | 125.1 | 1 | <0.0001 |
| | | | Time | 32.46 | 2 | <0.0001 |
| | | | Interaction | 15.36 | 2 | <0.0001 |
| Figure 4D (80 nmol/kg biMOG$_{Low}$:monoTfR:BACE1 vs 160 nmol/kg monoTfR:BACE1) | | | | | | |
| | *Two-way ANOVA (only datapoints until day 7 included)* | 0.05 | Treatment | 50.82 | 1 | <0.0001 |
| | | | Time | 95.33 | 2 | <0.0001 |
| | | | Interaction | 10.9 | 2 | 0.0001 |
| Figure 4E (160 nmol/kg biMOG$_{Low}$:monoTfR:BACE1 vs 160 nmol/kg monoTfR:BACE1) | | | | | | |
| | *Two-way ANOVA (only datapoints until day 7 included)* | | Treatment | 18.02 | 1 | 0.0004 |
| | | | Time | 24.20 | 2 | <0.0001 |
| | | | Interaction | 4.682 | 2 | 0.0208 |
| Figure 4E (80 nmol/kg biMOG$_{Low}$:monoTfR:BACE1 vs 160 nmol/kg monoTfR:BACE1) | | | | | | |
| | *Two-way ANOVA (only datapoints until day 7 included)* | 0.05 | Treatment | 8.762 | 1 | 0.0077 |
| | | | Time | 41.96 | 2 | <0.0001 |
| | | | Interaction | 3.754 | 2 | 0.0413 |
| Figure 5B | | | | | | |
| | *Two-way ANOVA* | 0.05 | Treatment | 0.02315 | 1 | 0.8802 |
| | | | Time | 102.3 | 3 | <0.0001 |
| | | | Interaction | 0.6036 | 3 | 0.6181 |
| Figure 5C | | | | | | |
| | *Two-way ANOVA* | 0.05 | Treatment | 1.107 | 1 | 0.3016 |
| | | | Time | 2.461 | 3 | 0.0833 |

| | | | | | | |
|---|---|---|---|---|---|---|
| | | | Interaction | 0.5864 | 3 | 0.6289 |
| Figure 5D | | | | | | |
| | *Two-way ANOVA* | 0.05 | Treatment | 0.4625 | 1 | 0.5020 |
| | | | Time | 16.13 | 3 | <0.0001 |
| | | | Interaction | 9.884 | 3 | 0.0001 |
| Figure 5E | | | | | | |
| | *Two-way ANOVA* | 0.05 | Treatment | 4.617 | 1 | 0.0404 |
| | | | Time | 0.5711 | 3 | 0.6387 |
| | | | Interaction | 1.062 | 3 | 0.3810 |
| Figure 8C (Brain) | | | | | | |
| | *Two-way ANOVA (only datapoints until day 7 included)* | 0.05 | Treatment | 1593 | 1 | <0.0001 |
| | | | Time | 1394 | 3 | <0.0001 |
| | | | Interaction | 254.5 | 3 | <0.0001 |
| Figure 8C (Spinal Cord) | | | | | | |
| | *Two-way ANOVA (only datapoints until day 7 included)* | 0.05 | Treatment | 202.7 | 1 | <0.0001 |
| | | | Time | 136.9 | 3 | <0.0001 |
| | | | Interaction | 23.23 | 3 | <0.0001 |
| Figure 9A (monoTfR:BACE1) | | | | | | |
| | *One-way ANOVA* | 0.01 | Time | 0.6872 | 3 | 0.5822 |
| Figure 9A (biMOG$_{Low}$:monoTfR:BACE1) | | | | | | |
| | *One-way ANOVA* | 0.01 | Time | 2.082 | 3 | 0.1729 |
| Figure 9A (biMOG$_{Low}$:monoTfR:SarsCov) | | | | | | |
| | *One-way ANOVA* | 0.01 | Time | 13.94 | 3 | 0.0010 |
| Figure 9B (monoTfR:BACE1) | | | | | | |
| | *One-way ANOVA* | 0.01 | Time | 1.481 | 3 | 0.2844 |
| Figure 9B (biMOG$_{Low}$:monoTfR:BACE1) | | | | | | |
| | *One-way ANOVA* | 0.01 | Time | 14.45 | 3 | 0.0009 |
| Figure 9B (biMOG$_{Low}$:monoTfR:SarsCov) | | | | | | |
| | *One-way ANOVA* | 0.01 | Time | 4.090 | 3 | 0.0436 |
| Figure 9C (monoTfR:BACE1) | | | | | | |
| | *One-way ANOVA* | 0.01 | Time | 36.92 | 3 | <0.0001 |
| Figure 9C (biMOG$_{Low}$:monoTfR:BACE1) | | | | | | |
| | *One-way ANOVA* | 0.01 | Time | 5.067 | 3 | 0.0251 |
| Figure 9C (biMOG$_{Low}$:monoTfR:SarsCov) | | | | | | |
| | *One-way ANOVA* | 0.01 | Time | 7.166 | 3 | 0.0093 |
| Supplementary Figure 2 | | | | | | |
| | *One-way ANOVA* | 0.05 | Treatment | 2392 | 6 | <0.0001 |

*Table S4. Pairwise comparisons from ANOVAs of supplemental table 3.*

| Figure | Groups | Mean Difference | df | Adjusted P value |
|---|---|---|---|---|
| Figure 3A | D1 noMOG - biMOG$_{High}$ | -21.88 | 30 | 0.5740 |
| | D1 noMOG-monoMOG$_{High}$ | -27.95 | 30 | 0.3641 |
| | D1 noMOG-biMOG$_{Low}$ | -3.422 | 30 | 0.9991 |
| | D1 noMOG-monoMOG$_{Low}$ | -2.265 | 30 | 0.9998 |
| | D3 noMOG-biMOG$_{High}$ | 5.425 | 30 | 0.9945 |
| | D3 noMOG-monoMOG$_{High}$ | 1.795 | 30 | >0.9999 |
| | D3 noMOG-biMOG$_{Low}$ | -0.8450 | 30 | >0.9999 |
| | D3 noMOG-monoMOG$_{Low}$ | -0.4425 | 30 | >0.9999 |
| Figure 3B | D1 PBS-biMOG$_{High}$ | 39.49 | 31 | <0.0001 |
| | D1 PBS-monoMOG$_{High}$ | 52.27 | 31 | <0.0001 |
| | D1 PBS-biMOG$_{Low}$ | 45.70 | 31 | <0.0001 |
| | D1 PBS-monoMOG$_{Low}$ | 47.11 | 31 | <0.0001 |
| | D1 PBS-noMOG | 38.49 | 31 | <0.0001 |
| | D3 PBS-biMOG$_{High}$ | 37.47 | 31 | 0.0017 |
| | D3 PBS-monoMOG$_{High}$ | 36.92 | 31 | 0.0020 |
| | D3 PBS-biMOG$_{Low}$ | 39.69 | 31 | 0.0009 |
| | D3 PBS-monoMOG$_{Low}$ | 42.11 | 31 | 0.0005 |
| | D3 PBS-noMOG | 32.16 | 31 | 0.0071 |
| Figure 3C | D1 noMOG - biMOG$_{High}$ | 4.945 | 30 | 0.4769 |
| | D1 noMOG-monoMOG$_{High}$ | -4.965 | 30 | 0.4734 |
| | D1 noMOG-biMOG$_{Low}$ | -2.753 | 30 | 0.8659 |
| | D1 noMOG-monoMOG$_{Low}$ | -2.465 | 30 | 0.9038 |
| | D3 noMOG-biMOG$_{High}$ | -20.83 | 30 | <0.0001 |
| | D3 noMOG-monoMOG$_{High}$ | -13.99 | 30 | 0.0023 |
| | D3 noMOG-biMOG$_{Low}$ | -34.21 | 30 | <0.0001 |
| | D3 noMOG-monoMOG$_{Low}$ | -7.285 | 30 | 0.1689 |
| Figure 3D | D1 PBS-biMOG$_{High}$ | 25.65 | 31 | <0.0001 |
| | D1 PBS-monoMOG$_{High}$ | 34.62 | 31 | <0.0001 |
| | D1 PBS-biMOG$_{Low}$ | 30.23 | 31 | <0.0001 |
| | D1 PBS-monoMOG$_{Low}$ | 40.12 | 31 | <0.0001 |
| | D1 PBS-noMOG | 36.36 | 31 | <0.0001 |
| | D3 PBS-biMOG$_{High}$ | 31.92 | 31 | 0.0001 |
| | D3 PBS-monoMOG$_{High}$ | 37.76 | 31 | <0.0001 |
| | D3 PBS-biMOG$_{Low}$ | 37.57 | 31 | <0.0001 |
| | D3 PBS-monoMOG$_{Low}$ | 37.56 | 31 | <0.0001 |
| | D3 PBS-noMOG | 14.94 | 31 | 0.0831 |
| Figure 9A (monoTfR:BACE1) | PBS-D1 | -0.04514 | 9 | 0.8998 |
| | PBS-D3 | -0.02028 | 9 | 0.9888 |
| | PBS-D7 | -0.1094 | 9 | 0.4275 |
| Figure 9A (biMOG$_{Low}$:monoTfR:BACE1) | PBS-D1 | -0.1029 | 9 | 0.6877 |
| | PBS-D3 | -0.07422 | 9 | 0.8443 |
| | PBS-D7 | -0.2672 | 9 | 0.0890 |
| Figure 9A (biMOG$_{Low}$:monoTfR:SarsCov) | PBS-D1 | -0.2024 | 9 | 0.0547 |
| | PBS-D3 | 0.2741 | 9 | 0.0119 |
| | PBS-D7 | 0.1496 | 9 | 0.1684 |
| Figure 9B (monoTfR:BACE1) | PBS-D1 | 0.001096 | 9 | >0.9999 |

| | | | | |
|---|---|---|---|---|
| | PBS-D3 | 0.1998 | 9 | 0.2165 |
| | PBS-D7 | 0.02771 | 9 | 0.9882 |
| Figure 9B (biMOG$_{Low}$:monoTfR:BACE1) | PBS-D1 | -0.05137 | 9 | 0.9795 |
| | PBS-D3 | -0.3504 | 9 | 0.4130 |
| | PBS-D7 | -0.9856 | 9 | 0.0005 |
| Figure 9B (biMOG$_{Low}$:monoTfR:SarsCov) | PBS-D1 | 0.08263 | 9 | 0.6737 |
| | PBS-D3 | 0.2224 | 9 | 0.0704 |
| | PBS-D7 | 0.2619 | 9 | 0.0339 |
| Figure 9C (monoTfR:BACE1) | PBS-D1 | 0.1493 | 9 | 0.0184 |
| | PBS-D3 | 0.4501 | 9 | <0.0001 |
| | PBS-D7 | 0.1841 | 9 | 0.0055 |
| Figure 9C (biMOG$_{Low}$:monoTfR:BACE1) | PBS-D1 | 0.2023 | 9 | 0.1078 |
| | PBS-D3 | -0.03687 | 9 | 0.9531 |
| | PBS-D7 | -0.1502 | 9 | 0.2657 |
| Figure 9C (biMOG$_{Low}$:monoTfR:SarsCov) | PBS-D1 | 0.03286 | 9 | 0.9913 |
| | PBS-D3 | -0.1415 | 9 | 0.6473 |
| | PBS-D7 | -0.5698 | 9 | 0.0073 |
| Supplementary Figure 2 | monoGFP:BACE1-biMOG$_{Low}$:monoTfR:SarsCov | -2286 | 23 | <0.0001 |
| | monoGFP:BACE1-biMOG$_{Low}$:monoTfR:BACE1 | -515.0 | 23 | <0.0001 |
| | monoGFP:BACE1-monoTfR:SarsCov | -122.0 | 23 | 0.0005 |
| | monoGFP:BACE1-monoTfR:BACE1 | -54.80 | 23 | 0.2143 |
| | monoGFP:BACE1-biMOG$_{Low}$:monoGFP:SarsCov | -127.0 | 23 | 0.0003 |
| | monoGFP:BACE1-biMOG$_{Low}$:monoGFP:BACE1 | -10.68 | 23 | 0.9926 |

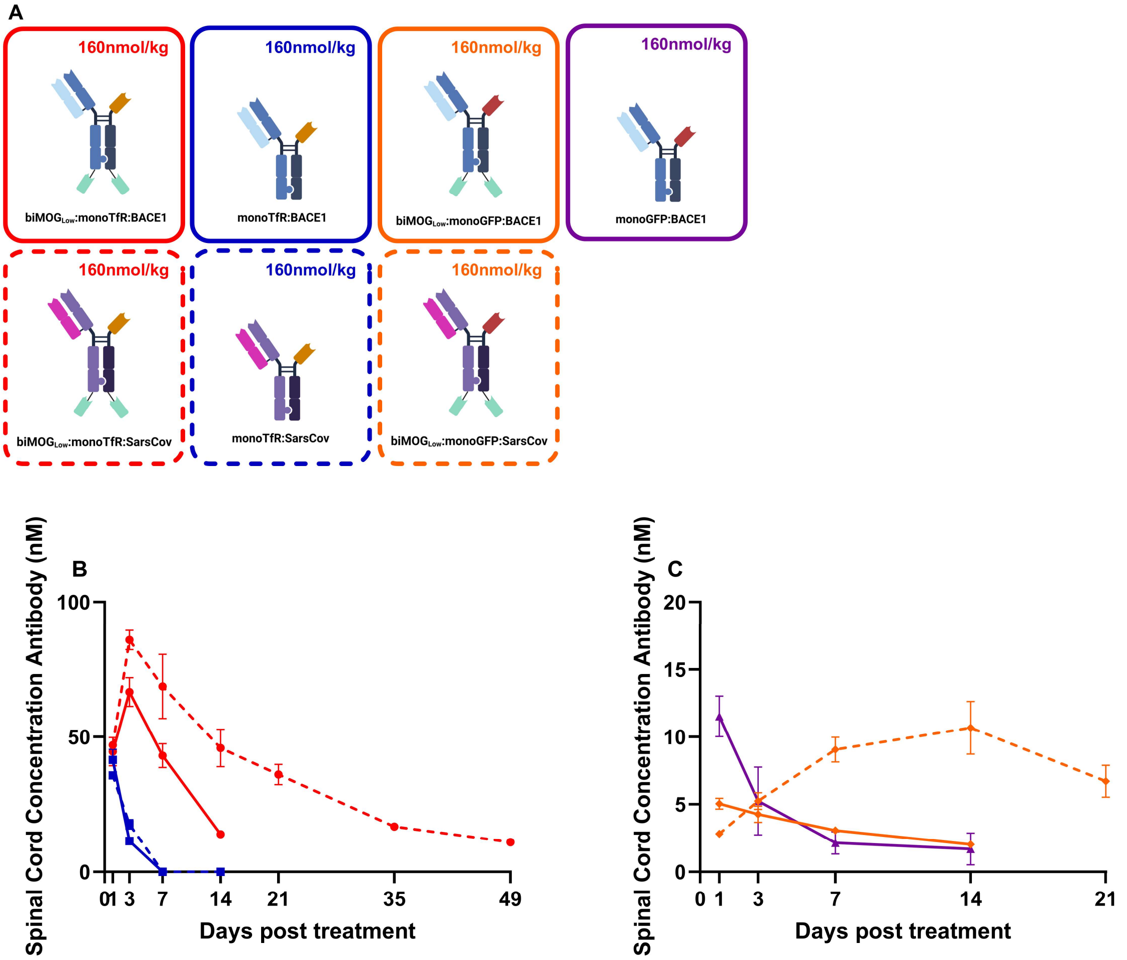


***Figure S1. Spinal cord antibody concentrations.*** *Mice received a single intravenous injection of 160 nmol/kg of monoTfR:BACE1/SarsCov (blue/dashed blue), biMOG$_{Low}$:monoTfR:BACE1/SarsCov (red/dashed red) or biMOG$_{Low}$:monoGFP:BACE1/SarsCov (orange/dashed orange). PK was evaluated over a period of 7-49 days.* ***(A)*** *Schematic representation of the antibodies .* ***(B)*** *Spinal cord antibody levels of actively shuttled antibodies.* ***(C)*** *Spinal cord antibody levels of antibodies without TfR binding. Curves represent mean ± SEM (n = 3-6 per group).*

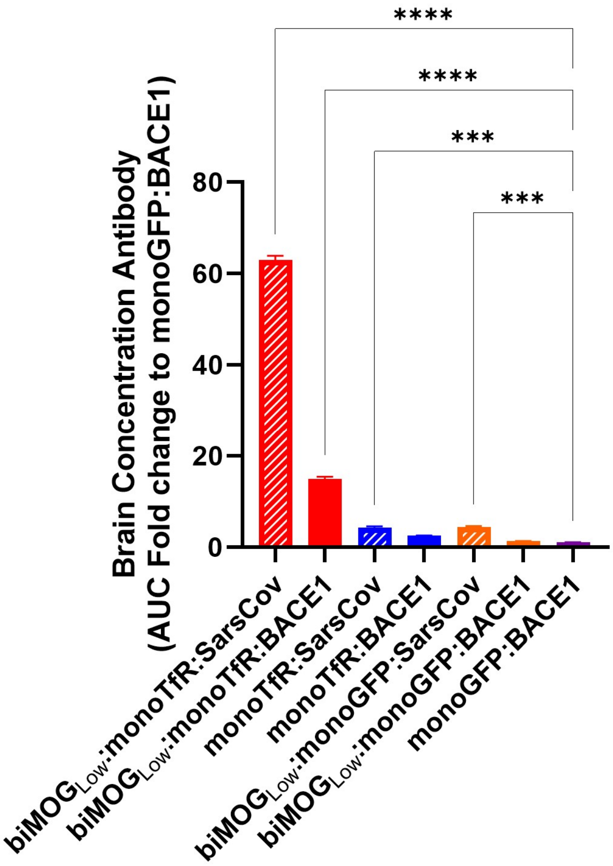


***Figure S2. AUC of antibody brain concentrations.*** *Area under the curve of brain concentration of antibodies after mice received a single IV dose of 160 nmol/kg. Bar graphs represent mean ± SEM (n = 3-6 per group). Statistical test: one-way ANOVA with Dunnett's multiple comparisons test compared to control treatment monoGFP:BACE1. (*p<0.05, **p<0.01, ***p<0.001, ****p<0.0001)*

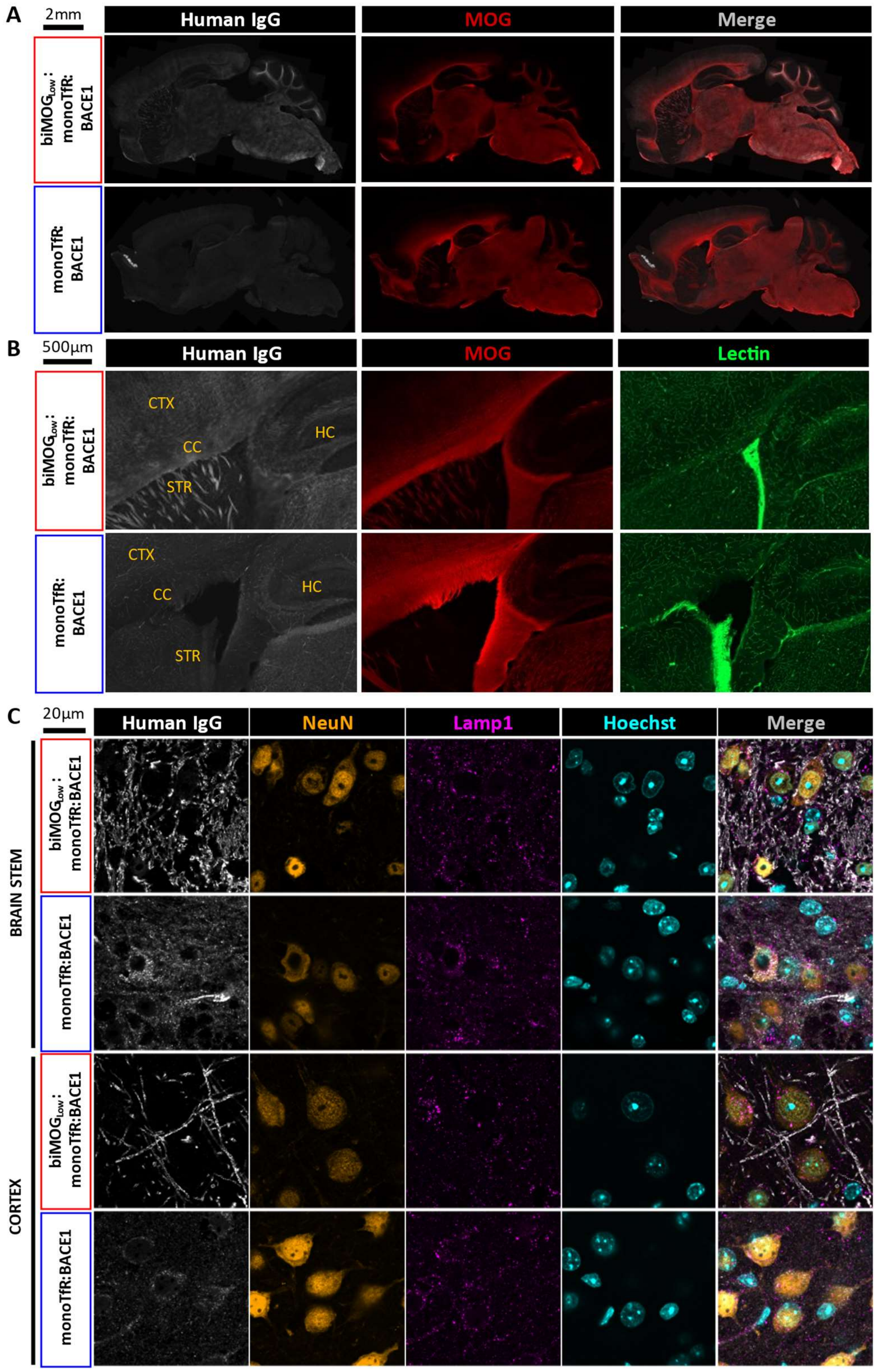


***Figure S3. Human IgG localization in mouse brains.*** *Immunohistochemistry of mouse brain sagittal sections 3 days post single 160 nmol/kg IV dose of biMOG$_{Low}$:monoTfR:BACE1 or monoTfR:BACE1. Representative images are shown from n=3 animals/group, n=2 IHC sections/animal.* ***(A)*** *Low magnification imaging of mouse brain sagittal sections immunostained for human IgG (white) and MOG (red) showing distinct distribution profiles.* ***(B)*** *Zoom of overview scans in A, displaying part of the cortex (CTX), corpus callosum (CC), hippocampus (HC) and striatum (STR) stained for human IgG (white), MOG (red) and lectin (green), demonstrate a more vascular staining pattern of monoTfR:BACE1 compared to biMOG$_{Low}$:monoTfR:BACE1.* ***(C)*** *Confocal imaging of cortex and brain stem, immunostained for human IgG (white), NeuN (orange), Lamp1 (Magenta) and Hoechst (Cyan) indicating more neuronal internalization of monoTfR:BACE1 compared to biMOG$_{Low}$:monoTfR:BACE1.*

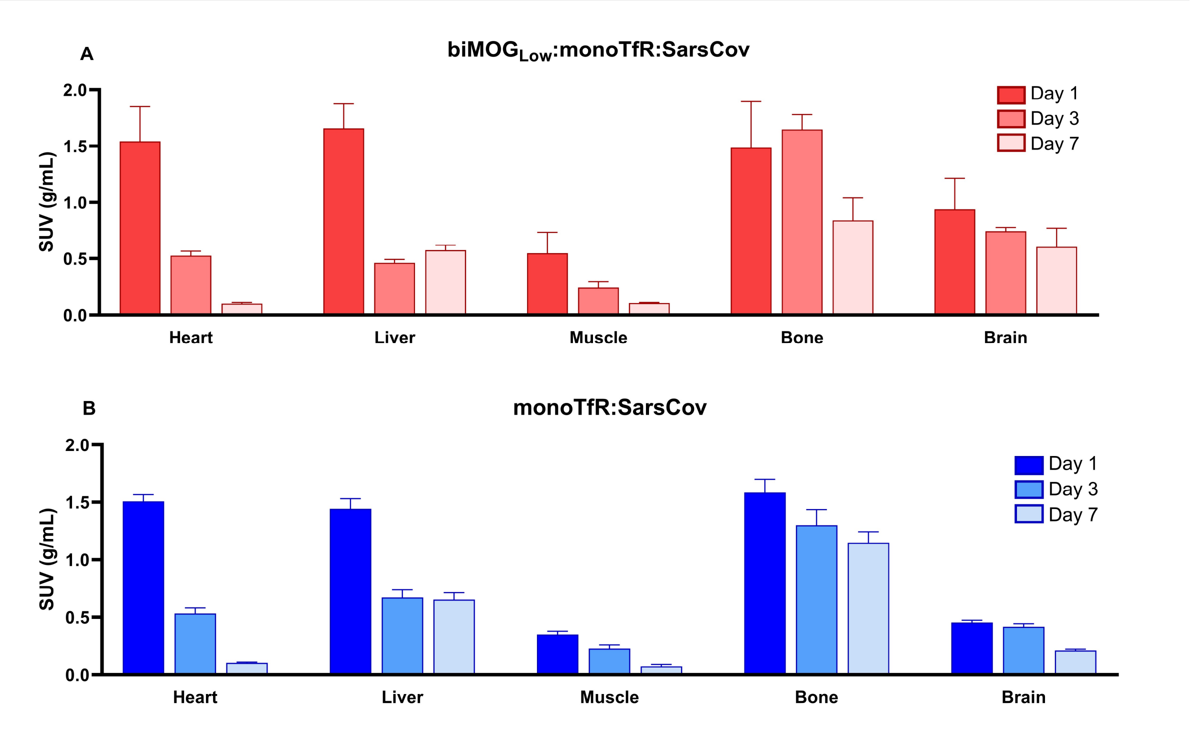


***Figure S4. In vivo biodistribution of [$^{89}$Zr]Zr-DFO*-biMOG$_{Low}$:monoTfR:SarsCov and [$^{89}$Zr]Zr-DFO*-monoTfR:SarsCov.*** *Mice received a single IV injection of* ***(A)*** *[$^{89}$Zr]Zr-DFO*-biMOG$_{Low}$:monoTfR:BACE1 and* ***(B)*** *[$^{89}$Zr]Zr-DFO*-monoTfR:BACE1. µPET imaging was performed to quantify radiolabeled antibodies in different organs. Bar graphs represent mean standardized uptake value (g/mL) ± SEM (n = 2-4 per group).*
